# Plasticity of the binding pocket in peptide transporters underpins promiscuous substrate recognition

**DOI:** 10.1101/2023.02.14.528348

**Authors:** Vadim Kotov, Maxime Killer, Katharina E. J. Jungnickel, Jian Lei, Giada Finocchio, Josi Steinke, Kim Bartels, Jan Strauss, Florine Dupeux, Anne-Sophie Humm, Irina Cornaciu, José A. Márquez, Els Pardon, Jan Steyaert, Christian Löw

## Abstract

Proton-coupled oligopeptide transporters (POTs) are promiscuous transporters of the Major Facilitator Superfamily, that constitute the main route of entry for a wide range of dietary peptides and orally administrated peptidomimetic drugs. Given their clinical and pathophysiological relevance, several bacterial and mammalian POT homologs have been extensively studied on a structural and molecular level. However, the molecular basis of recognition and transport of the wide range of peptide substrates has remained elusive. Here we present 14 X-ray structures of the bacterial POT DtpB in complex with chemically diverse di- and tripeptides, providing novel insights into the plasticity of the conserved central binding cavity. We analyzed binding affinities for more than 80 peptides and monitored uptake by a fluorescence-based transport assay. To probe if all natural 8400 di- and tripeptides can bind to DtpB, we employed state-of-the-art molecular docking and machine learning and conclude that peptides of a specific subset with compact hydrophobic residues are the best DtpB binders.

## Introduction

Living cells have to adapt rapidly to environmental changes to maintain nutrient homeostasis, which requires the use of different nitrogen-containing nutrients. Hence, cells express a variety of genes to ensure the scavenging of alternative nitrogen sources such as amino acids or peptides ^1^. Peptide transporters of the major facilitator superfamily (MFS), namely the proton-dependent oligopeptide transporter (POT) family, provide the cell with valuable nitrogen and carbon sources by mediating the uptake of di- and tripeptides ^2^. POT members are known to have an overall helical arrangement termed “MFS-fold”, formed by two helical bundles (N- and C-terminal bundle) that are related by a pseudo-two-fold symmetry ^3–8^ and function according to the alternate access mechanism ^9–13^. Additionally, they carry specific sequence signature motifs important for proton coupling, ligand binding, and transport ^2,3,14^. They are known to be highly promiscuous, expected to transport almost all 8400 di- and tripeptides composed of proteinogenic amino acids ^2^.

In *E. coli*, four members of the POT family were identified and termed di- and tripeptide permease (Dtp) A to D ^15–17^. Experimental structures of DtpA, DtpC and DtpD were reported recently ^7,8,18^. Transport inhibition experiments indicated that the binding sites of DtpA and DtpB interact with a large number of substrates and peptidomimetic drugs in a similar fashion as the extensively studied mammalian homolog PepT1 ^16,19–24^. DtpC and DtpD, however, were classified as atypical POTs, as they were shown previously to favour positively charged peptides as substrates ^18,22,25–30^.

In the last decade, over 50 entries of the POT family were deposited in the Protein Data Bank (PDB), representing eleven bacterial and two mammalian homologs, bound to eight unique natural di- and tripeptides (Ala-Phe, Ala-Glu, Ala-Gln, Ala-Leu, Phe-Ala, Phe-Ala-Gln, Ala-Ala-Ala, alafosfalin), peptidomimetic drugs (valaciclovir, valganciclovir, 5-aminolevulinic acid) and a potent transport inhibitor Lys[Z(NO2)]-Val ^3–8,10,18,31–41^. These efforts revealed the basic principle underlying peptide binding in POTs which can be described as an electrostatic clamping mechanism between the invariable part of peptides (N- and C-termini as well as the peptide backbone) and conserved residues in the transporter (mainly via arginine, lysine, glutamate, asparagine and tyrosine residues). In addition, the molecular changes during transport have been recently described in human PepT1 and PepT2, providing insights into the dynamics of peptide transporters and the local rearrangements of the binding site through-out the full transport cycle ^31^. However, it still remains unclear how POTs can recognize and transport such a vast variety of peptides due to the lack of high-resolution structures of POTs bound to chemically diverse substrates.

Here, we determined crystal structures of DtpB complexed with 14 different di- and tripeptides, providing novel insights into the plasticity of the conserved central binding cavity in response to a wide range of chemically diverse peptides. We thereby also complete the entire family of experimental POT structures from *E. coli* ranging from DtpA to D. Moreover, we measured binding affinities for more than 80 peptides using the thermal shift method and employed a fluorescence-based transport assay to monitor the uptake of peptides into liposomes reconstituted with DtpB. Our analysis indicates that high affinity peptides are only poorly transported and rather act as inhibitors, while peptides in the medium affinity range display the highest transport rates. Finally, we employed state-of-the-art molecular docking and machine learning to probe if all 8400 di- and tripeptides composed of proteinogenic amino acids can bind to DtpB and conclude that a specific subset of peptides with compact hydrophobic residues are the best DtpB binders.

## Results and Discussion

### Structures of DtpB-peptide complexes stabilized by nanobody 132

In order to obtain highly diffracting crystals of DtpB, we selected conformation-specific nanobodies after immunizing a noninbred llama ^42^. Out of 31 recombinantly expressed and purified nanobodies, 14 bound DtpB with a dissociation constant of 30 nM or lower as evident by bilayer interferometry (BLI) measurements (Figure 1, Supplementary Figure 1, Supplementary Tables 1 and 2). They were used as crystallization chaperones in subsequent crystallization trials and nanobody 132 (Nb132) emerged as the most promising binder for co-crystallization approaches. DtpB-Nb132 was initially incubated with the tripeptide Ala-Leu-Ala (ALA), and well-diffracting crystals grew using the vapor diffusion method. The structure of DtpB was determined using the atomic model of DtpA ^7^ (PDB accession number 6GS4) as search model for molecular replacement. Strong positive peaks in the difference electron density map (i.e. F_obs_-F_calc_) indicated the presence of additional electrons/atoms in the periplasmic region of DtpB, and within the central cavity, allowing modelling of Nb132 and the tripeptide.

**Figure 1:**
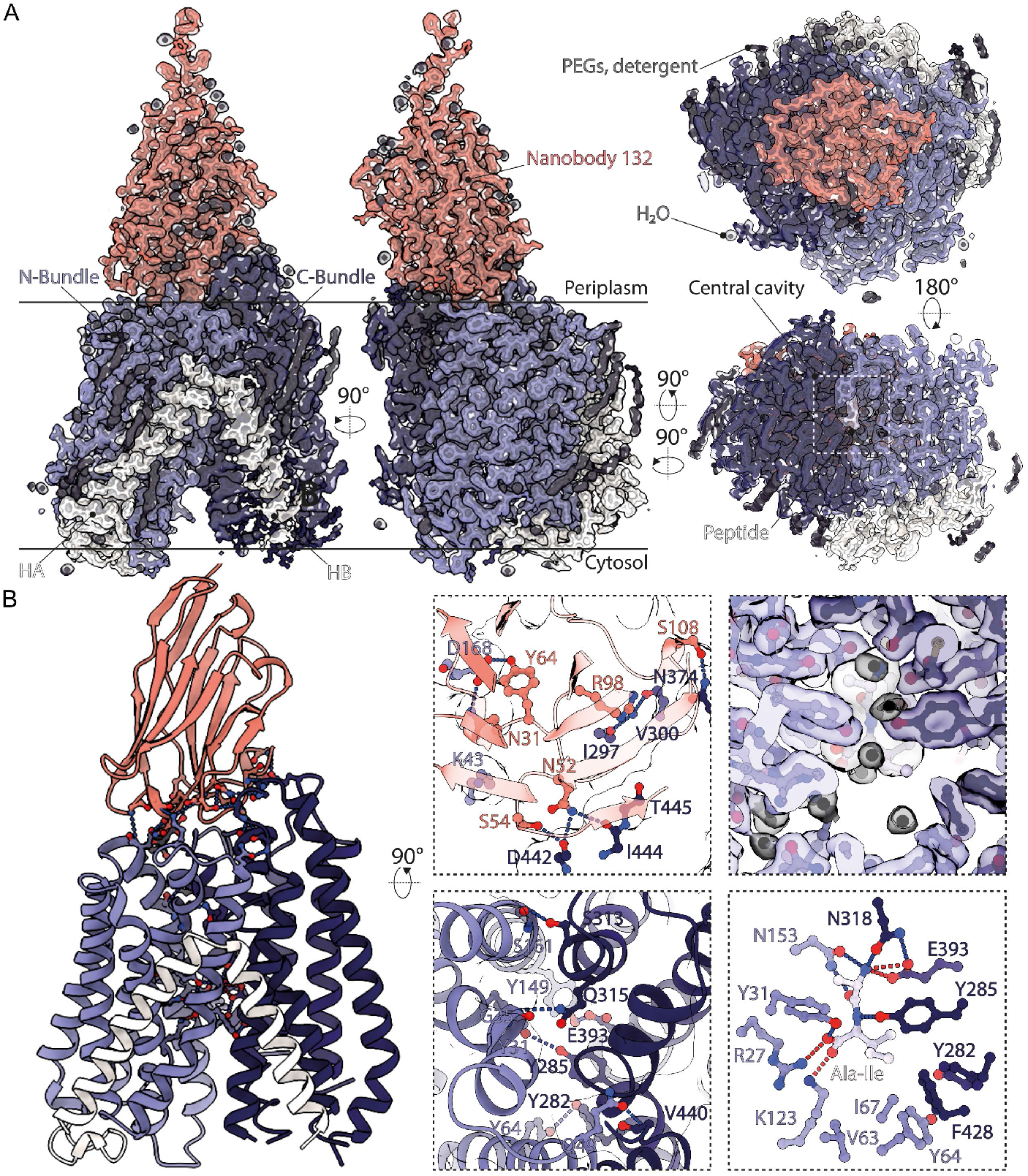
Structure of DtpB bound to nanobody Nb132 and the dipeptide AI. (A) The atomic model of DtpB-Nb132 fitted from the highest resolution dataset AI. The 2Fo-Fc map is shown as transparent surface (at σ = 1). The different structural elements are labeled. (B) Residues stabilizing the observed conformation are displayed as ball and sticks and the secondary structural elements are shown as ribbons. Interactions between the transporter and the nanobody are shown in the top left close up view, while interactions stabilizing the IF state between both bundles are highlighted in the bottom panel. The electron density at the binding site and the dipeptide are illustrated in the top right close up view, and the electrostatic interactions between the peptide and DtpB are shown as dashes in the bottom panel (red dashed lines denote salt bridges; blue dashed lines correspond to polar interactions; waters are shown in black).

DtpB crystallized in an inward facing open (IF open) state. In this conformation the central cavity of the transporter is open to the cytoplasm and closed to the periplasm (Figure 1). DtpB adopts the canonical MFS fold, characterized by two six transmembrane helix bundles. The N- and C-terminal bundles are linked by the HA-HB helices, as observed in other bacterial POTs ^3–8,10,18,32–35,37,39^. The overall structure of DtpB is similar to DtpA, DtpC, and DtpD with C_α_ atom RMSD (root-mean-square deviation) values of 0.9, 1.3, and 1.2 Å, respectively. In DtpB, the IF open state is stabilised by electrostatic interactions between the two bundles i.e., Y31-Y285; G35-Q315; Q49-V440; Y64-Y282; and Y149-E393. Nb132 further stabilizes the closure on the periplasmic side by interacting with the periplasmic surface of the transporter through polar contacts (Figure 1B).

In order to obtain a broader vision of the plasticity of the binding site in response to peptides possessing various chemical groups, the co-crystallization efforts were continued with an in-house library of 82 peptides. Several batches of the DtpB-Nb132 complex were prepared, incubated with a peptide, and then dispensed robotically in 96 well screens containing three sets of crystallization conditions. If the diffraction resolution of an obtained crystal was better than 4 Å, the chemical screens were further refined around the best conditions. In this campaign, X-ray diffraction data were collected and analyzed for more than 2000 crystals with the help of automated crystallography pipelines based on the CrystalDirect technology ^43,44^. For each dataset, with a resolution limit higher than 3.5 Å, the presence or absence of a peptide was initially assessed by using an atomic model of DtpB devoid of any substrate, to calculate a difference electron density map (i.e., F_obs_-F_calc_). There, strong positive peaks within the central cavity indicated the presence of a ligand. The co-crystallized peptide was then modeled inside the positive peak, as a di- or trialanine moiety first, and then mutated to its original sequence, as the signal improved during the refinement steps. In the final validation, OMIT maps excluding the modeled ligands, were calculated for each structure (Supplementary Figure 2). In summary, we obtained 14 unique peptide bound datasets in a resolution range between 2.0 and 2.8 Å with the following peptides: Ala-Phe (AF), Ala-Ile (AI), Ala-Leu (AL), Ala-Gln (AQ), Ala-Val (AV), Ala-Trp (AW), Lys-Val (KV), Met-Ser (MS), Asn-Val (NV), Ser-Leu (SL), Ala-Leu-Ala (ALA), Ala-Phe-Ala (AFA), Ala-Pro-Phe (APF) and Ala-Trp-Ala (AWA) (Figure 2, Supplementary Figure 3, Table 1). This represents a large portfolio of di- and tripeptides for POTs compared to the present literature data. These structures now allowed us to analyze changes and local rearrangements in the binding site of DtpB and shed light on how promiscuity is achieved in this transporter family.

**Figure 2:**
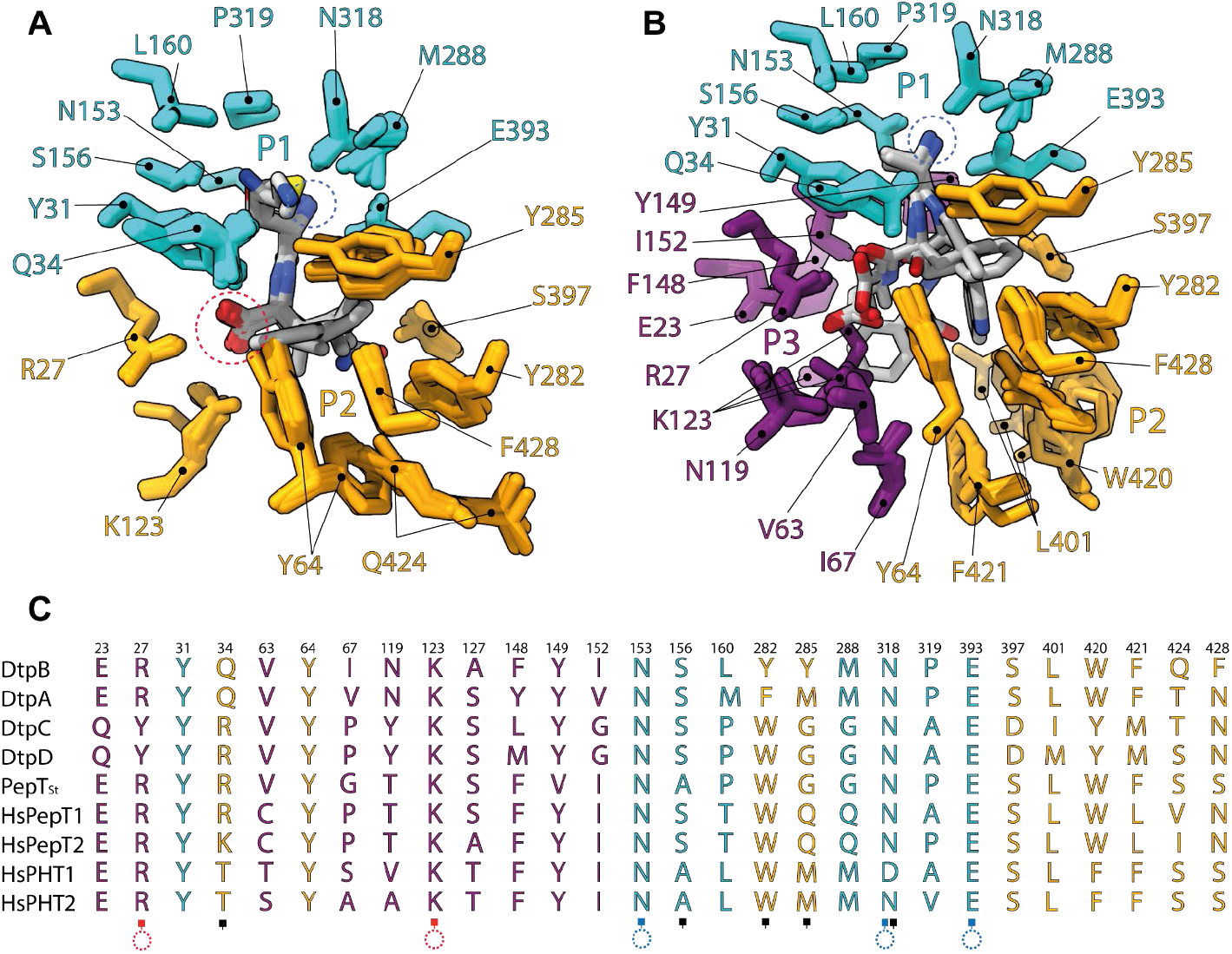
Definition of the binding pocket of DtpB. (A) Superimposition of the dipeptide co-crystal structures. (B) Superimposition of the tripeptide co-crystal structures. (C) Sequence alignment of the residues constituting the P1, P2, and P3 pockets in POTs. Residues constituting the P1, P2, and P3 pockets are respectively colored in cyan, yellow, and purple. Blue and red dashed rings circle the N-termini, and the C-termini. Blue and red squares indicate residues mediating electrostatic interactions with the termini of the co-crystallized peptides. Black squares indicate residues mediating polar interactions with the side chains of co-crystallized peptides. The N-termini are all coordinated in the same manner, while the C-termini adopt different positions in tripeptides.

**Table 1:** Data collection and refinement statistics of DtpB-Nb132-peptide datasets.

| Peptide | AF | AFA | AI | AL | ALA | APF | AQ | AV | AW | AWA | KV | MS | NV | SL |
| --- | --- | --- | --- | --- | --- | --- | --- | --- | --- | --- | --- | --- | --- | --- |
| PDB code | 8B18 | 8B19 | 8B1A | 8B1B | 8B1C | 8B1D | 8B1E | 8B1F | 8B1G | 8B17 | 8B1H | 8B1I | 8B1K | 8B1J |
| Resolution (Å) | 49.90 - 2.30<br>(2.38 - 2.30) | 70.30 - 2.45<br>(2.55 - 2.45) | 51.69 - 2.15<br>(2.23 - 2.15) | 51.95 - 2.80<br>(2.90 - 2.80) | 51.79 - 2.56<br>(2.65 - 2.56) | 62.58 - 2.30<br>(2.38 - 2.30) | 58.74 - 2.47<br>(2.56 - 2.47) | 84.47 - 2.35<br>(2.43 - 2.35) | 59.16 - 2.50<br>(2.59 - 2.50) | 58.76 - 2.50<br>(2.59 - 2.50) | 69.95 - 2.60<br>(2.69 - 2.60) | 50.95 - 2.55<br>(2.64 - 2.55) | 54.14 - 2.80<br>(2.90 - 2.80) | 51.16 - 2.67<br>(2.77 - 2.67) |
| Space group | P 2 21 21 | P 2 21 21 | P 2 21 21 | P 2 21 21 | P 2 21 21 | P 2 21 21 | P 2 21 21 | P 2 21 21 | P 2 21 21 | P 2 21 21 | P 2 21 21 | P 2 21 21 | P 2 21 21 | P 2 21 21 |
| Cell dimensions<br>a, b, c (Å) | 54.42, 125.07,<br>168.71 | 54.70, 125.51,<br>169.72 | 54.32, 125.74,<br>168.00 | 54.59, 125.84,<br>169.03 | 54.40, 123.54,<br>169.33 | 54.76, 125.16,<br>169.90 | 54.47, 125.39,<br>168.18 | 54.65, 126.17,<br>168.94 | 54.61, 126.41,<br>168.04 | 54.60, 125.36,<br>168.82 | 54.34, 125.71,<br>168.37 | 54.18, 124.68,<br>167.46 | 54.14, 124.93,<br>165.77 | 54.49, 125.66,<br>168.04 |
| α, β, γ (°) | 90, 90, 90 | 90, 90, 90 | 90, 90, 90 | 90, 90, 90 | 90, 90, 90 | 90, 90, 90 | 90, 90, 90 | 90, 90, 90 | 90, 90, 90 | 90, 90, 90 | 90, 90, 90 | 90, 90, 90 | 90, 90, 90 | 90, 90, 90 |
| Total No. of reflections | 667300<br>(64221) | 578325<br>(55600) | 810303<br>(71235) | 378285<br>(38108) | 75320 (7396) | 105697<br>(10434) | 83033 (8176) | 649577<br>(45053) | 81675 (7972) | 80967 (8074) | 72749 (7106) | 75797 (7414) | 57042 (5622) | 65106 (5161) |
| No. of reflections | 52149 (5144) | 43863 (4296) | 63584 (6245) | 29527 (2894) | 37673 (3699) | 52854 (5217) | 42145 (4129) | 49598 (4863) | 40878 (4000) | 40497 (4037) | 36376 (3553) | 37899 (3707) | 28523 (2811) | 32661 (2628) |
| Completeness (%) | 99.93 (99.94) | 99.90 (99.95) | 99.94 (99.94) | 99.88 (100.00) | 99.89 (99.78) | 99.93 (99.96) | 99.71 (99.61) | 99.88 (99.94) | 99.15 (98.62) | 98.65 (100.00) | 99.92 (100.00) | 99.92 (99.95) | 99.77 (99.93) | 96.74 (77.81) |
| I/σI | 16.79 (1.47) | 16.45 (1.40) | 18.87 (1.48) | 16.30 (1.57) | 16.74 (1.89) | 13.91 (1.24) | 14.56 (1.82) | 15.18 (0.78) | 12.36 (1.05) | 14.29 (1.43) | 13.06 (1.04) | 9.44 (1.30) | 12.01 (0.88) | 12.66 (1.19) |
| Wilson B-factor (Å²) | 53.52 | 71.01 | 50.5 | 80.22 | 66.52 | 57.56 | 61.28 | 59.41 | 69.68 | 69.65 | 71.69 | 54.22 | 77.16 | 80.58 |
| R<sub>merge</sub> | 0.08998<br>(1.649) | 0.09384<br>(2.310) | 0.07757<br>(1.664) | 0.13730<br>(2.112) | 0.02248<br>(0.4068) | 0.02381<br>(0.5902) | 0.02899<br>(0.4735) | 0.11130<br>(3.0180) | 0.01985<br>(0.6883) | 0.02077<br>(0.4964) | 0.02634<br>(0.6764) | 0.04223<br>(0.5812) | 0.03269<br>(0.8924) | 0.02147<br>(0.675) |
| R<sub>meas</sub> | 0.09399<br>(1.719) | 0.09789<br>(2.405) | 0.08102<br>(1.742) | 0.14350<br>(2.198) | 0.03179<br>(0.5752) | 0.03368<br>(0.8347) | 0.0410<br>(0.6697) | 0.11610<br>(3.199) | 0.02807<br>(0.9734) | 0.02937<br>(0.702) | 0.03725<br>(0.9565) | 0.05973<br>(0.822) | 0.04623<br>(1.262) | 0.03037<br>(0.9545) |
| R<sub>pim</sub> | 0.02673<br>(0.4834) | 0.02738<br>(0.6643) | 0.02302<br>(0.5110) | 0.04087<br>(0.6025) | 0.02248<br>(0.4068) | 0.02381<br>(0.5902) | 0.02899<br>(0.4735) | 0.03238<br>(1.0420) | 0.01985<br>(0.6883) | 0.02077<br>(0.4964) | 0.02634<br>(0.6764) | 0.04223<br>(0.5812) | 0.03269<br>(0.8924) | 0.02147<br>(0.6750) |
| CC1/2 | 0.999 (0.648) | 0.999 (0.522) | 0.999 (0.591) | 0.997 (0.531) | 0.999 (0.694) | 1 (0.609) | 0.998 (0.640) | 0.991 (0.285) | 1 (0.432) | 1 (0.643) | 1 (0.488) | 1 (0.545) | 1 (0.367) | 1 (0.350) |
| CC* | 1 (0.887) | 1 (0.828) | 1 (0.862) | 0.999 (0.833) | 1 (0.905) | 1 (0.870) | 1 (0.883) | 0.998 (0.666) | 1 (0.777) | 1 (0.885) | 1 (0.810) | 1 (0.840) | 1 (0.733) | 1 (0.720) |
| Reflections used in refinement | 52127 (5142) | 43856 (4295) | 63564 (6244) | 29508 (2894) | 37667 (3699) | 52830 (5216) | 42133 (4122) | 49579 (4862) | 40846 (3993) | 40479 (4037) | 36360 (3553) | 37882 (3705) | 28489 (2809) | 32590 (2577) |
| Reflections used for R<sub>free</sub> | 2589 (251) | 2223 (236) | 3113 (316) | 2999 (306) | 1892 (188) | 2648 (249) | 2078 (213) | 2514 (252) | 2015 (191) | 2000 (193) | 1807 (180) | 1838 (200) | 1382 (128) | 1621 (122) |
| R<sub>work</sub> | 0.2267<br>(0.3774) | 0.2241<br>(0.3115) | 0.2154<br>(0.3490) | 0.2254<br>(0.3354) | 0.2151<br>(0.2944) | 0.2299<br>(0.3515) | 0.2221<br>(0.3108) | 0.2164<br>(0.2998) | 0.2245<br>(0.3442) | 0.2314<br>(0.3054) | 0.2257<br>(0.3659) | 0.2241<br>(0.3319) | 0.2260<br>(0.3869) | 0.2211<br>(0.3589) |
| R<sub>free</sub> | 0.2536<br>(0.4101) | 0.2673<br>(0.3354) | 0.2390<br>(0.3854) | 0.2633<br>(0.3708) | 0.2410<br>(0.3293) | 0.2575<br>(0.3866) | 0.2669<br>(0.3498) | 0.2363<br>(0.3244) | 0.2481<br>(0.3548) | 0.2647<br>(0.3278) | 0.2602<br>(0.3766) | 0.2574<br>(0.3686) | 0.2503<br>(0.4129) | 0.2636<br>(0.3964) |
| Number of non-hydrogen atoms | 4718 | 4793 | 4787 | 4669 | 4781 | 4792 | 4786 | 4785 | 4786 | 4791 | 4787 | 4784 | 4789 | 4787 |
| Protein residues | 580 | 581 | 580 | 580 | 581 | 581 | 580 | 580 | 580 | 581 | 580 | 580 | 580 | 580 |
| R.m.s. deviations |  |  |  |  |  |  |  |  |  |  |  |  |  |  |
| Bond lengths (Å) | 0.011 | 0.011 | 0.01 | 0.012 | 0.012 | 0.01 | 0.011 | 0.01 | 0.01 | 0.012 | 0.012 | 0.011 | 0.013 | 0.01 |
| R.m.s. deviations |  |  |  |  |  |  |  |  |  |  |  |  |  |  |
| Angles (°) | 1.32 | 1.3 | 1.08 | 1.44 | 1.45 | 1.32 | 1.32 | 1.1 | 1.32 | 1.32 | 1.37 | 1.3 | 1.56 | 1.19 |
| Ramachandran favored (%) | 98.42 | 97.72 | 98.06 | 96.83 | 98.42 | 98.42 | 98.42 | 98.06 | 98.06 | 97.54 | 97.89 | 97.18 | 97.89 | 97.36 |
| Ramachandran allowed (%) | 1.41 | 2.11 | 1.94 | 3.17 | 1.58 | 1.58 | 1.58 | 1.94 | 1.94 | 2.46 | 2.11 | 2.82 | 1.94 | 2.64 |
| Ramachandran outliers (%) | 0.18 | 0.18 | 0 | 0 | 0 | 0 | 0 | 0 | 0 | 0 | 0 | 0 | 0.18 | 0 |
| Rotamer outliers (%) | 0.42 | 1.06 | 1.06 | 1.91 | 0.64 | 0.64 | 0.64 | 1.06 | 1.7 | 0.85 | 0.64 | 0.85 | 0.64 | 1.06 |
| Clashscore | 10.1 | 10.09 | 8.1 | 10.94 | 10.3 | 8.61 | 7.05 | 7.57 | 9.15 | 9.24 | 9.46 | 9.15 | 11.36 | 10.83 |
| Average B-factor | 67.63 | 80.49 | 60 | 82.94 | 75.22 | 67.08 | 64.85 | 67.4 | 77.25 | 75.84 | 75.99 | 61.75 | 82.31 | 81.74 |

Statistics for the highest-resolution shell are shown in parentheses.

### Plasticity in the peptide binding pocket to accommodate diverse peptides

All DtpB complexes crystallized in the same space group with very similar cell dimensions (Table 1). A superposition of the structures highlights that the N-termini of all di- and tripeptides are anchored in a similar manner (Figure 2). The primary amine is steadily hooked by N153, N318, and E393 of DtpB. This triad of residues remains fixed in all structures, and is conserved in all prototypical POTs, with the exception of the peptide-histidine transporter 1 (PHT1), where N318 is replaced by an aspartate. Additional residues of the binding site form a pocket around the N-terminal residue of the substrate (Supplementary Figures 3 and 4). These include Y31, Q34, S156, S156, L160, M288, and P319. Together with the conserved N153, N318, E393 triad, they constitute the so-called P1 pocket. The chemical diversity of N-terminal residues in the co-crystallized tripeptide data sets was poor (only alanine residues), but richer for dipeptides (i.e., alanine, lysine, serine, methionine, asparagine). P1 does not undergo conformational changes in presence of these various residues, however, polar interactions stabilize certain substrates (Supplementary Figure 3, Figure 2). Notably, Q34 interacts with the ε-amino group of K1* in KV; N318 with the thioester group of M1* in MS; S156 and N318 with the hydroxymethyl group of S1* in SL; and S156, and N318 with the carboxamide group of N1* in NV. The C-terminus of dipeptides adopted a constant position, stabilized by R27 and occasionally by K123 as well.

The picture is different for the second residue of the substrate. To adapt the central binding cavity to the different sizes of side chains carried by the C-terminal residues of dipeptides (P2 pocket), two rotamer conformations of Y64 and Q424 are possible (Supplementary Figure 5). They control the volume of the ‘upper region’ of the P2 pocket. For instance, in presence of the small dipeptide AV, the upper region of P2 is tightened by the conformation of Y64 and Q424, while it is widened in the presence of the bulky dipeptide AW (Supplementary Figure 5). Additional residues such as Y285 and F428 further fine-tune the upper region of P2. Of note, Y285 and Y64 delimit the P2 pocket from the P1 and P3 pockets respectively (Figure 2). Unlike other side-chains fitting in P2, the indole moiety of W2* in the tripeptide AWA extends further down, towards the cytosolic side of the C-bundle (Supplementary Figure 5). This ‘lower region’ of P2 (i.e. L401, W420, and F421) is more flexible, and closes up to stabilize W2*. Polar interactions also occur between P2 and the peptides. Y282 stabilizes the indole ring of W2* in AWA as well as the carboxamide group of Q2* in AQ (Supplementary Figure 3 F,J). In addition, the hydrophobic side chains in the second position (i.e. in the peptides AV, AL, AI, AF, AW, ALA, AFA, AWA) are increasingly stabilized as a function of their size, and through contraction of the upper region of P2.

The backbone coordination of tripeptides withstands different torsion angles around amide bonds. Since the primary amine of the N-terminus remains hooked in place between N153, N318, and E393, the carboxylic group of the C-terminus is subsequently shifted or rotated, resulting in different poses (Supplementary Figure 3, Figure 2). For instance, the carboxylic group of AWA coincides with the ones of dipeptides. This co-localization is achieved by a kinked backbone geometry of the tripeptide. AFA and ALA are not kinked, but stretched. In comparison, in dipeptides AL and AF a conformational change in K123 creates sufficient space for the carboxylic group of L2* and A3*, to fit in a position preserving the stabilizing salt bridges with R27 and K123 (Supplementary Figures 3 and 6). Finally, in the case of the tripeptide APF, the proline residue restrains the backbone and evicts the C-terminus from the positively charged patch formed by R27 and K123, and the bulky phenyl group of F3* extends towards the cytosolic side of the N-bundle, in P3 (Supplementary Figures 3 and 6). The overall plasticity of the binding site of DtpB is illustrated in Movie 1.

In summary, the N153, N318, E393 triad is a common anchor point of peptides N-termini, in agreement with earlier literature ^32,45,46^. We find that the C-termini of peptides are often stabilized by R27 and K123, but the latter is not mandatory, contrary to the previously suggested model ^14^. Recent molecular dynamics (MD) studies using human PepT2 and the tripeptide AAA ^36^ support this observation and suggest that peptides engage with the binding pocket via A1* first, before being tightly locked in place by the triad (N192, N348, E622 in human PepT2). The simulations also indicate that R27 and K64 later contribute to further stabilization of the C-terminus. Importantly, K64 (Q34 in DtpB) was not essential in DtpB to coordinate the tripeptide APF and K64 is generally not conserved among POT homologs. It is likely that the presence of bulky side chains in the N-terminal position, would cause local changes in P1, but we did not succeed in determining structures of such complexes. The versatility of P2 was previously described with rearrangements of Y68 and W427 in a POT from *Streptococcus thermophilus* (PepT_St_) ^32,33^, which correspond to Y64 and W420 in DtpB. Here, these two residues critically contribute to adapt P2 to the various co-crystallized peptides, but other residues (Y282, Y285, L401, Q424, F428) are also involved. Except for the different rotamer conformations of K123 to enable various positions of tripeptides, the P3 pocket was rather stable compared to P2. Although these observations indicate that the plasticity of POTs originates mainly in the P2 pocket, many more combinations of peptides are able to bind to and be transported by DtpB and other prototypical POTs. Besides, not all residues involved in ligand promiscuity are conserved in POTs. This could explain the differences of substrate affinities between homologs reported in the literature.

### Determination of peptide affinities using a thermal unfolding assay

To shed further light on the interactions of different di- and tripeptides with DtpB and determine binding affinities for a large set of peptides, we employed a thermal unfolding assay. This approach is commonly used to characterize the stability of proteins and their functional interactions ^47–50^. In particular, the stability of proteins (as judged by the melting temperature T_m_) increases in a concentration-dependent manner when a ligand is added (Figure 3A). Various concepts were proposed to obtain a ligand dissociation constant (K_D_) based on the ligand-induced shifts of T_m_ ^51,52^. These approaches typically assume classic thermodynamic behavior of proteins during unfolding, i.e. the equilibrium is quickly reached at all temperatures, and protein unfolding is fully reversible. The latter is rarely observed in practice, so a kinetic description of the unfolding process should be used instead ^53,54^. Recently Hall introduced and validated a kinetic model to determine affinities from thermal shifts (Figure 3B) ^55^. We modified Hall’s model to take advantage of modern non-linear curve fitting methods (see Materials & Methods) and applied it to DtpB titrated with di- and tripeptides. Titration curves could be fit with high confidence for peptides in a broad affinity range (Figure 3C). The resulting K_D_ was not strongly affected by assay conditions ^56^ (Supplementary Figure 7A). Importantly, our approach allows us to predict K_D_ of new peptides obtained from T_m_ measurements at a single concentration (see Materials & Methods, Supplementary Figure 7B). The rank order of the K_D_ agreed well with an orthogonal technique (microscale thermophoresis (=MST), Supplementary Figure 7C). With this approach we could efficiently measure K_D_ values for 82 di- and tripeptides (Figure 3D) to DtpB and cover a broad region of the peptide chemical space. As expected, determined affinities span several orders of magnitude, with the tightest binders showing affinities in the low μM range while others interacted with DtpB only poorly or not at all.

**Figure 3:**
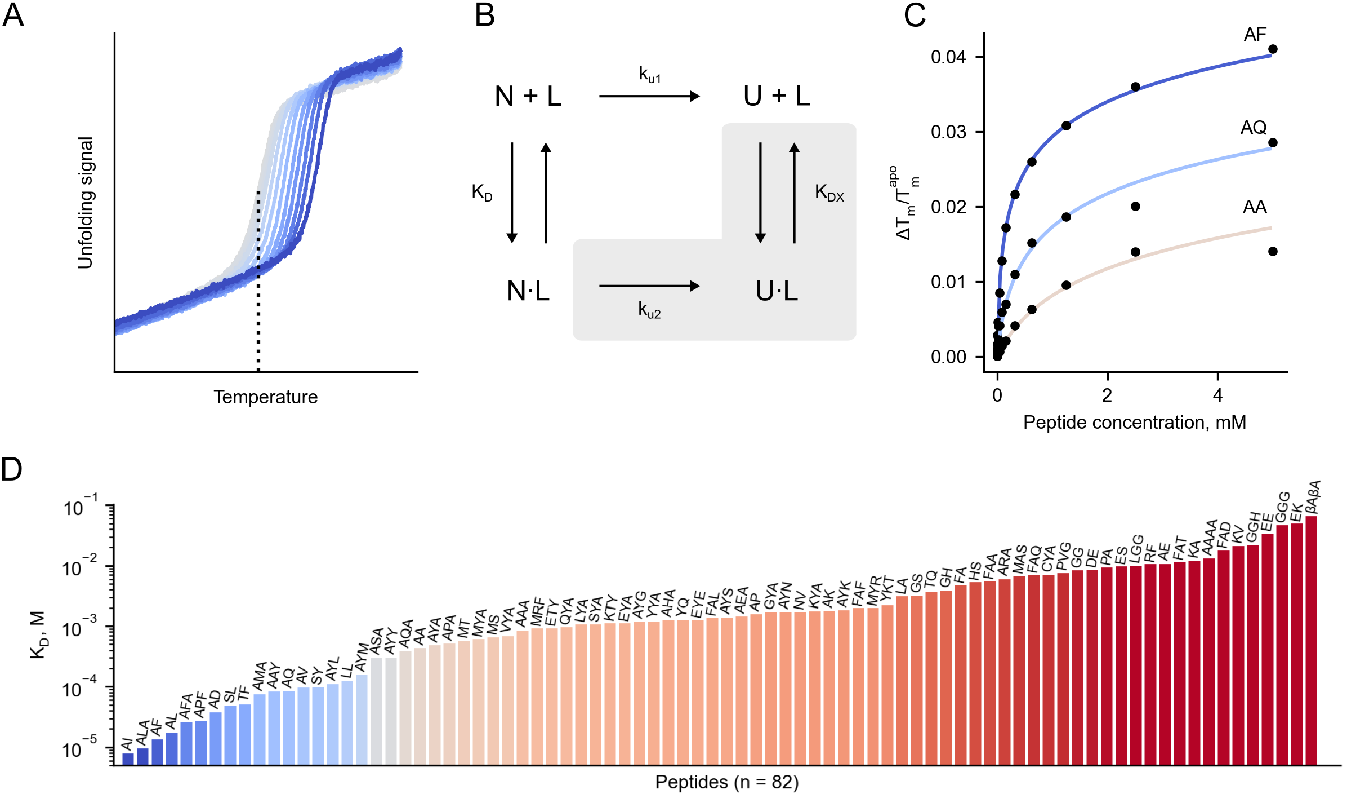
Large-scale determination of peptide binding to DtpB using a thermal unfolding assay. (A) Thermal unfolding assays can quantify protein-ligand interactions; ligand concentration increases with the blue color intensity. Melting temperature (T_m_) of the apo state (gray curves) is shown with a vertical dashed line. (B) Hall’s irreversible unfolding model to describe receptor-ligand interaction in a thermal unfolding assay. N - native state of the receptor, U - unfolded state of the receptor, L - ligand. Elements under the gray area denote the part of the model that was omitted in the current study. (C) Exemplary saturation curves of diverse peptides. Peptide affinity is color-coded with AF having the highest affinity, and AA having the lowest affinity. T_m_^apo^ is the melting temperature of the apo state, ΔT_m_ is the difference in T_m_ between the apo state and in presence of respective peptide concentration. (D) Overview of K_D_ values obtained in this study; the affinity is color-coded with low-affinity peptides in red and high-affinity peptides in blue.

### Tight peptide binding is not associated with transport

The core biological function of a transporter is its ability to move molecules across the lipid bilayer. To gain more detailed insights into the uptake of various peptides via DtpB, we established a robust transport assay in liposomes, termed ‘pyranine assay’. Since the peptide uptake by POTs is coupled to protons ^57–59^ we used the pH sensitive fluorescent dye pyranine ^60^ to indirectly monitor peptide transport into DtpB-containing liposomes. Such an approach was recently applied to characterize the bacterial POT PepT_St_ ^61^. We utilized similar experimental conditions to follow transport activity of DtpB reconstituted into liposomes (Figure 4A). To validate the assay, we initially confirmed transport of the known POT substrates Ala-Ala (AA) and Gly-Gly (GG) by DtpB (Figure 4B) and related transporter PepT_St_ ^16,61^ (data not shown).

**Figure 4:**
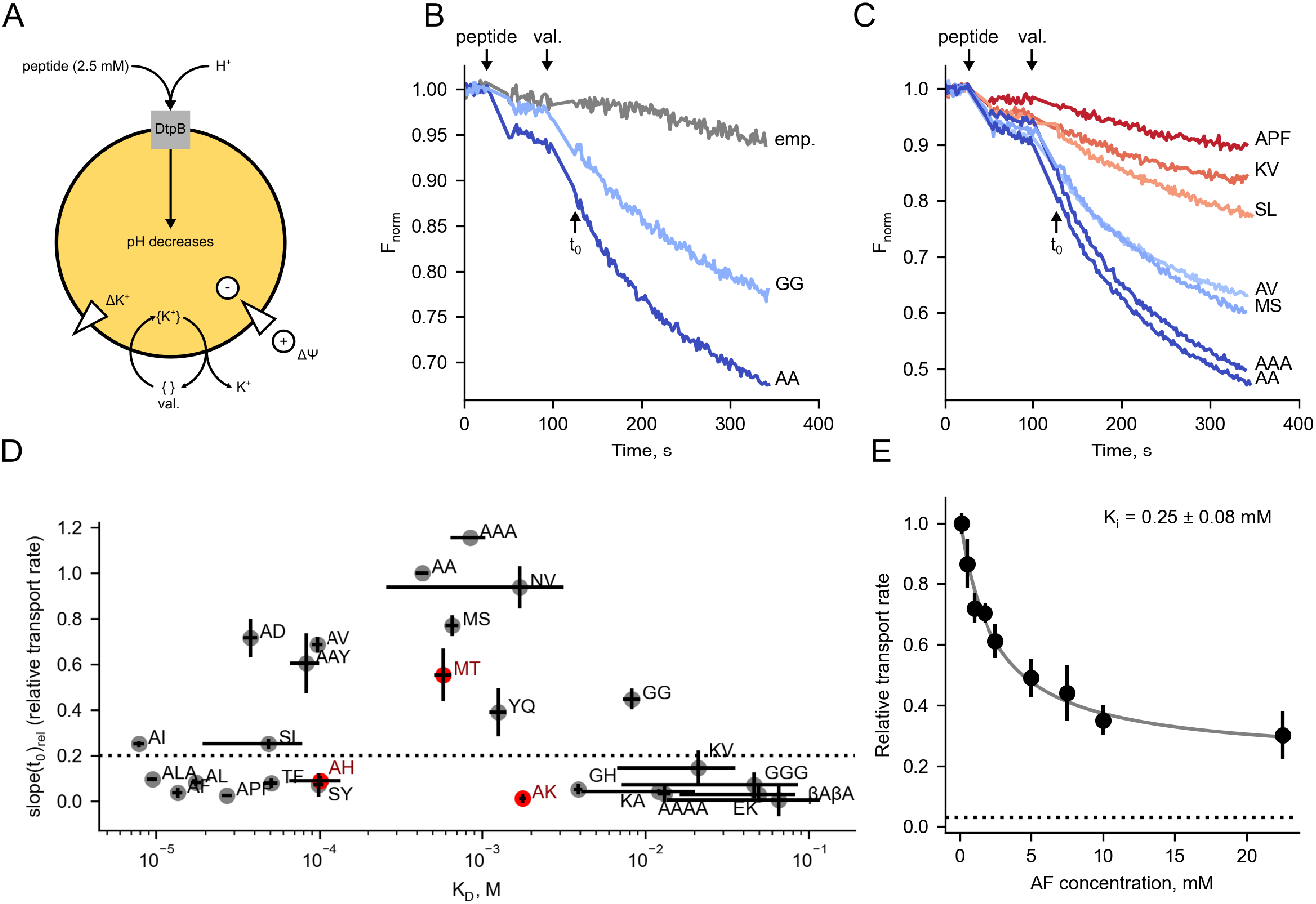
Measurement of peptide transport by DtpB and its relationship with peptide K_D_. (A) The principle of the pyranine assay. DtpB is reconstituted into liposomes, and a concentration gradient of potassium ions (ΔK^+^) from the inside to the outside of the liposome is created. Upon addition of valinomycin (val.) the potassium ions are chelated ({K^+^}) and carried across the membrane and establish an electrochemical gradient (membrane potential ΔΨ). The membrane becomes hyperpolarized, and DtpB can utilize this gradient for proton-coupled peptide transport (peptide is added at indicated concentration to the outside buffer only). Proton flux into the liposome changes the fluorescence spectrum of the membrane-impermeable dye pyranine (yellow) present inside the liposome. (B) Exemplary transport curves of AA and GG detected with the pyranine assay. The transport signal from liposomes without DtpB (emp.) is shown in gray. Upper arrows denote approximate time when the peptide or valinomycin (val.) were added. Addition of peptide and valinomycin requires the fluorescence reading to be paused. Another arrow indicates t_0_, which corresponds to the time point when the fluorescence readings are resumed. (C) Exemplary transport curves of diverse peptides tested in this work with pyranine assay. Approximate time when the peptide or valinomycin (val.) were added is shown with arrows. The curves are color-coded to show slow or no transport in red, and fast transport in blue. (D) Relationship between binding (K_D_) and slope(t_0_)_rel_ (relative initial transport rate in pyranine assay). Dashed horizontal line indicates a cut-off of 0.20, which corresponds to 20% of slope(t_0_) of AA. Only peptides above the dashed line are considered to be transported. Peptides marked in red were predicted to be transported based on their K_D_ and subsequently tested in the pyranine assay. Error bars for K_D_ are standard deviations (n = 2). Error bars for transport rate are the median absolute deviation (n = 2). (E) AF inhibits transport of AA. Relative transport rates of AA in presence of variable AF concentrations are shown as black dots, and the fit curve into Hill equation to determine IC_50_ is shown in grey. Error bars correspond to the standard deviation (n = 3). The relative transport rate of AF alone is shown with a dashed line.

To account for batch-to-batch variations between different liposome reconstitutions, we developed a non-linear curve fitting procedure to quantify the obtained transport curves in this assay (see Materials & Methods). In brief, the experimental data were corrected for the empty liposome signal and then fit to a single exponential decay function to obtain the time constant τ (tau) and the amplitude of the transport curve. The ratio of the amplitude to τ corresponds to the initial transport rate at time-point t_0_ (slope(t_0_)), corresponding to the time of the addition of valinomycin (Figure 4B). Transport measurements of the substrate AA and buffer alone served as a positive and a negative control and were used to normalize slope(t_0_) for all measured peptides to range from 0 to 1 thus obtaining slope(t_0_)_rel_. Quantification of transport rates at varying substrate concentrations allowed us to determine the apparent Michaelis-Menten constants (K_M_) for the dipeptides AA (0.29 ± 0.04 mM) and GG (2.63 ± 0.65 mM (Supplementary Figure 8A). These values are within the expected range for POTs ^10,40^. In addition, we used surface electrogenic event reader (SURFE^2^R) ^62^ as an orthogonal technique to verify the uptake of GG and AA by DtpB. In this assay an electrochemical gradient is absent, and the transport of peptides is driven by an excess of substrate in the external buffer. With this, we could confirm that AA and GG are transported by DtpB, however, the apparent K_M_ values were 20-30 fold higher compared to the electrochemical gradient driven conditions as used in the pyranine assay (Supplementary Figure 8).

Next, we determined slope(t_0_)_rel_ for 24 di- and tri-peptides using the pyranine assay, covering a broad spectrum of binding affinities as determined by thermal unfolding (Figure 4C,D). In the context of available K_D_ values, slope(t_0_)_rel_ forms a bell-shaped distribution (Figure 4D). Peptides that poorly bound to DtpB or not at all in our thermal shift assay exhibited low or no transport. This indicates that peptides with very low binding affinities would not initiate the transport cycle since they do not reside long or well enough bound in the binding site for any conformational changes of the transporter to occur. Alternatively, the correlation of low binding affinities and no apparent transport might also result from clashes or unfavoured positioning of the peptide in the binding site. The highest slope(t_0_)_rel_, i.e. peptides with at least 20% of the AA transport rate, could be observed for peptides with K_D_ values in the range of ~100 μM to ~2.5 mM. To confirm this observation, we measured transport for three more peptides picked from this affinity range: AH (K_D_ = 0.10 ± 0.04 mM), MT (K_D_ = 0.57 ± 0.07 mM) and AK (K_D_ = 1.77 ± 0.09 mM) (Figure 4D, red dots). Of those, only MT was transported suggesting that a K_D_ in a specific range is required, but not the only determining factor to enable transport by DtpB.

Our findings demonstrate that tightly bound peptides are either transported very slowly or act as inhibitors of transport. Interestingly, previous characterization of PepT_St_ using the pyranine assay ^61^ demonstrated transport of peptides AF and ALA, which are not transported by DtpB. To confirm the transport inhibitory effects of tightly binding peptides, we measured uptake of AA in presence of increasing concentrations of the high affinity binder AF (Figure 4E). The obtained IC_50_ value for AF is 2.37 ± 0.68 mM, which corresponds to an inhibitory constant K_i_ of 0.25 ± 0.08 mM. The IC_50_ value is two orders higher than the previously reported IC_50_ of ~ 0.027 mM for competition between radiolabeled AA and AF for PepT_St_ ^10^. This difference can be attributed to the different peptide concentrations used in the assay (2.5 mM of AA used in this work versus radiolabeled AA used at 0.03 mM concentration ^10^). We note that with our assay setup we did not aim to determine the number of protons being transported along with each peptide, though it was previously shown for PepT_St_ that this number may vary between di- and tripeptides ^61^. Consequently, the amplitude of the signal in the pyranine assay will be higher for peptides that carry more protons, while the time constant τ is not affected.

### DtpB preferentially binds small hydrophobic peptides

A more detailed analysis of the peptide K_D_ dataset (Figure 3G) indicates that DtpB preferentially binds peptides that are hydrophobic with a molecular weight below 300 Da (Supplementary Figure 9A). Since our dataset accounts for less than 1% of all possible di- and tripeptides made from proteinogenic amino acids (n = 8400), we asked whether the available crystal structures of DtpB-peptide complexes can be used to formulate more precise peptide recognition rules. For this, we performed flexible docking all of possible di- and tripeptides into DtpB by the Rosetta FlexPepDock protocol ^63^ and used a modified hit selection procedure (see Materials & Methods). This allowed us to predict the placement of the peptide inside DtpB for all 8400 di- and tripeptides (Figure 5A, Supplementary Figure 10).

**Figure 5:**
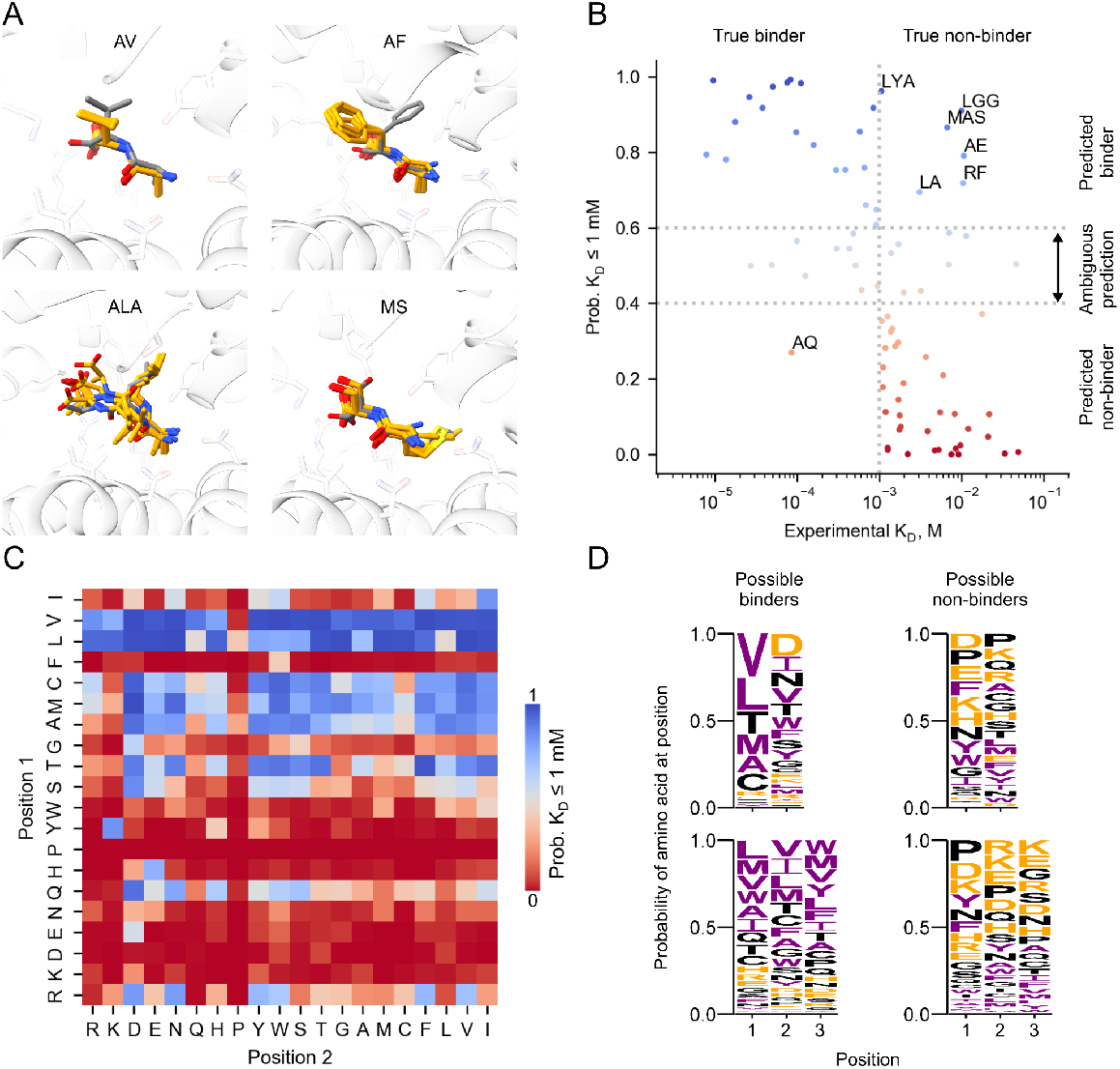
Flexible docking of di- and tripeptides into DtpB. (A) Exemplary docking results. The binding pocket is viewed from the cytoplasmic side, and only the native structure of DtpB is shown. The native conformation of the peptide is shown in gray, and top ten docked models are shown in orange. For all 14 peptides with known experimental structures see SupFigure 10. (B) Characterization of the logistic regression classifier that predicts binding of a peptide to DtpB. Vertical dashed line shows the value of experimental K_D_ that is used to divide peptides into two classes: binders (K_D_ ≤ 1 mM) and non-binders (K_D_ > 1 mM). Horizontal dashed lines correspond to the cut-off values for probabilities reported by the logistic regression classifier: if the binder probability is below 0.4, then it is very likely that the peptide will not bind to DtpB. On the other hand, with binder probabilities above 0.6 an interaction with DtpB is to be expected, however, it cannot be excluded that the peptide will still exhibit low affinity. In the probability range between 0.4 and 0.6 it is difficult to assign a peptide to any class (‘ambiguous’ prediction). Data points are colored by their binder probability (see color map in panel C). Labeled data points correspond to false-positives (top right quadrant) and false-negatives (lower left quadrant). (C) Influence of the amino acid identity on the probability of a dipeptide to be a DtpB binder. Amino acids in columns and rows are ordered by Kyte-Doolittle hydropathicity scale ^80^ with most hydrophobic residues in the top right corner. Note that in the Kyte-Doolittle scale W and Y are considered amphiphilic amino acids, so their hydropathicity is close to 0. (D) Sequence logos (consensus sequence representation) for dipeptides (top row) and tripeptides (bottom row) categorized by their probability to be a binder. Hydrophobic residues are shown in purple, charged residues are colored orange.

The Rosetta energy function is expressed in energy units (kcal/mol) and was recently calibrated to reliably represent real energies observed in protein molecules ^64^. On the other hand, it is impossible to account for all contributing factors that influence ligand binding without a thorough MD simulation. Thus, the scores reported by Rosetta or other docking applications generally show low or no correlation with experimentally observed affinity ^65,66^. Indeed, the rank order correlation (Spearman ρ) between the Rosetta score for docked peptides and their experimentally measured K_D_ was only −0.28 (data not shown). Therefore, we performed multiple linear regression by using individual Rosetta energy terms (e.g. electrostatic interactions, optimal placement of rotamers ^64^) as independent variables (features) and negative decimal logarithm of K_D_ as the dependent variable. The obtained models performed poorly in cross-validation (CV) tests with coefficient of determination (R^2^ score) close to 0 or negative (data not shown). Finally, we separated the di- and tripeptides with experimentally determined K_D_ into two classes: binders (K_D_ ≤ 1 mM) and non-binders (K_D_ > 1 mM) and trained a logistic regression classifier to distinguish these classes based on Rosetta energy terms (Figure 5B). Average area under curve (AUC) of the receiver-operator characteristic (ROC) during CV of this classifier was 0.67 ± 0.04 (see Materials & Methods), suggesting that it outperforms random classification (AUC 0.5). AUC of the classifier trained with full data was 0.88. The particular value of logistic regression is that it provides a probability estimate ^67^ for each sample to belong to each class (termed ‘binder probability’ in this work). If the peptide’s predicted binder probability is less than 0.4, then it is very likely to have an experimental K_D_ value above 1 mM (Figure 5B). On the other hand, several peptides with binder probability above 0.6 (AE, LA, LGG, LYA, MAS, RF) still may be poor binders when tested experimentally. This can be attributed to the fact that the solvent interaction is not modeled, which for instance plays an important role in peptide recognition by periplasmic binding proteins OppA ^68^ and DppA ^69^. Furthermore, ‘structural snapshots’ (as opposed to an MD simulation) are fundamentally limited in capturing the entropic component of binding ^62^. Still, the proposed classification approach allows us to exclude obvious non-binders (binder probability below 0.4) which constitute 4192 out of 8400 peptides (Supplementary Figure 9B). Only 727 peptides have binder probability over 0.99, i.e. in the search of new peptide binders to DtpB it is sufficient to experimentally check ~ 9% of all possible di- and tripeptides.

Next, we used the binder probability as a proxy to explore the peptide recognition landscape of DtpB (Figure 5C,D, Supplementary Figure 11). We observe critical importance of the first position in binding for both di- and tripeptides (Figure 5C, Supplementary Figure 11). In particular, polar amino acids but also bulky hydrophobic amino acids tend to decrease the binder probability (Figure 5C, Supplementary Figure 11). Our docking results also support the observation that DtpB preferentially binds hydrophobic peptides (Figure 5D), in agreement with previous data and an earlier study of the yeast homolog Ptr2p ^70^ and MD-based predictions for PepT_St_ ^71^. Interestingly, our analysis also reveals that tripeptide binders predominantly contain hydrophobic residues in position 2, whereas dipeptide binders may also contain hydrophilic residues in position 2 (Figure 5D).

### Prediction and validation of peptide pose and binding for DtpB

To make use of the above described assays and tools, we hypothesize that the combination of computational docking prediction tools and experiments can be integrated in a workflow to accelerate the identification of DtpB substrates (Figure 6A). Firstly, the computed docking poses can be used to estimate the binder probability for all possible di- and tripeptides. Secondly, top hits are validated with thermal unfolding assays to determine the binding affinity, which can be performed experimentally on a medium to high-throughput level. Lastly, the peptides that exhibit intermediate binding affinity (see the bell-shaped dependence of transport rate vs K_D_ in Figure 4D) are analyzed for uptake by a low-throughput transport assay.

**Figure 6:**
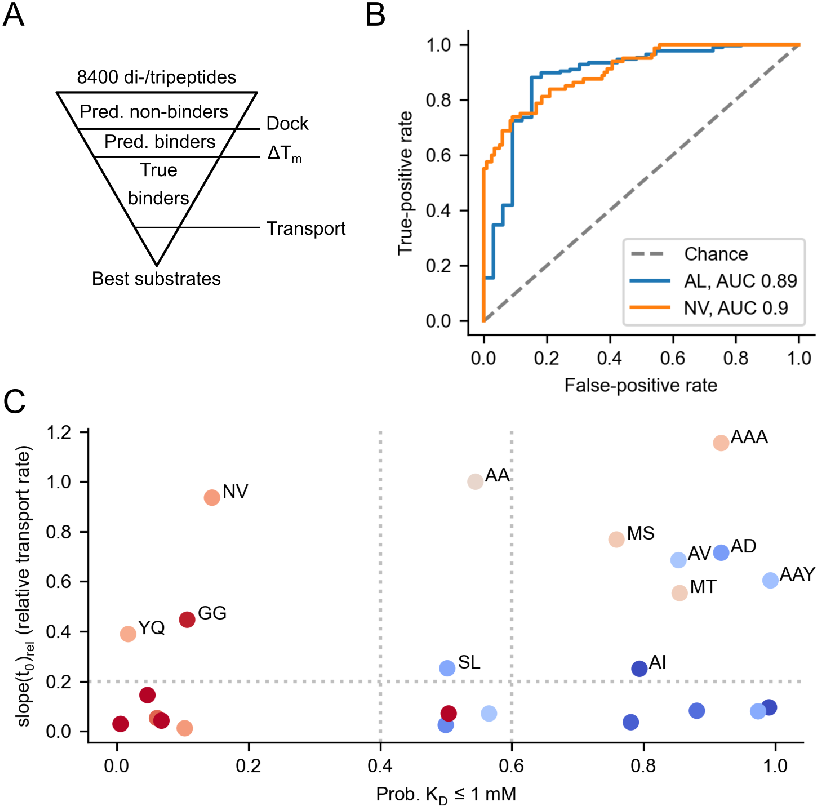
Characterization of the workflow to discover new peptide substrates for DtpB. (A) Schematic diagram of the proposed workflow. First, all possible di- and tripeptides (n=8400) composed of proteinogenic amino acids are separated into predicted binders and non-binders based on the docking results. This step is virtually instant, because the computation has already been performed. Second, true binders are identified using high-throughput K_D_ estimation based on thermal shifts (ΔT_m_). Finally, selected peptides in the optimal K_D_ range for transport (between ~100 μM and ~2.5 mM) are tested for transport using the low-throughput pyranine assay to identify true substrates. (B) ROC for the prediction of peptide docking poses for AL and NV. (C) Peptides transported by DtpB tend to have high binder probability. The shown data points are color-coded according to affinity: high affinity (i.e. low K_D_) in blue and low affinity (i.e. high K_D_) in red. Only peptides above the horizontal dashed line are considered transported. Vertical dashed lines indicate the ‘ambiguous probability’ region, i.e. where the binder/non-binder nature of the peptide is predicted with low confidence.

The obtained docking model (see DtpB preferentially binds small hydrophobic peptides) was developed with the input of twelve experimental structures of DtpB bound to various peptides determined in this study. The datasets of DtpB bound to dipeptides AL and NV have not been part of the training set and as such can be used to validate the predicted binding poses. RMSD values of the peptide backbone between the experimental structures and the top ten docking poses (rmsBB_offset see Materials & Methods) were 0.61 ± 0.16 Å for AL and 0.58 ± 0.15 Å for NV (Supplementary Figure 10). AUC of the modified hit selection procedure (when applied to all 200 generated dock poses) was 0.89 for AL and 0.9 for AV (Figure 6B), highlighting that the peptide binding pose inside the DtpB binding pocket can be predicted with high confidence. Furthermore, the RMSD for the peptide side-chains after superposition (rmsSC see Materials & Methods) was 1.45 ± 0.07 Å for AL and 1.25 ± 0.24 Å for NV, suggesting that the conformation of the peptide can be predicted with good precision.

To test whether the classifier between binder and non-binder peptides (see DtpB preferentially binds small hydrophobic peptides) can correctly predict binders, we experimentally measured the binding affinity of twelve commercially available peptides that were not used to train the classifier (Supplementary Table 3). Of those, five peptides (AH, AY, GD, LLA, RGD) had binder probabilities above 0.5 (predicted binders) and seven peptides (EA, GPE, HH, PY, SH, YA, YYR) had binder probabilities in the range of 0 - 0.1 (predicted non-binders). Five out of seven non-binders were true negatives (experimental K_D_ over 1 mM), and two predicted non-binders (PY and SH) turned out to be false negatives with K_D_ values of 206 ± 126 and 352 ± 38 μM. In case of predicted binders, three out of five (AH, AY, LLA) were true positives (experimental K_D_ below 1 mM). Two of the false-positives among predicted binders (RGD and GD) were in the ‘ambiguous’ probability region (Figure 5B) with binder probabilities of only 0.52 and 0.6 (indicating low confidence of the prediction). The ROC AUC of the experimental validation of the classifier is 0.74, i.e. the discrimination capability of our classification approach is acceptable, yet there is room for improvement.

Finally, we asked whether selection of peptides using binder probability could exclude the peptides that are known to be transported (see Tight peptide binding is not associated with transport). As demonstrated in Figure 6C, only three out of twelve peptides transported by DtpB have low binder probability (predicted non-binders). Two peptides, including the reference substrate AA, are in the ‘ambiguous’ probability region, so they are unlikely to be discovered with the proposed workflow, however, the remaining seven known substrates can be potentially identified starting from the docking analysis.

## Conclusions

In this work we present one of the most comprehensive structural and functional characterizations of a POT to date. We established a high-throughput crystallization pipeline, and determined the structures of 14 complexes of DtpB with ten different dipeptides and four tripeptides. Flexible residues within the binding site and multiple stabilizing polar interactions with peptides carrying various chemical groups were identified. Combined with a comprehensive biochemical characterization, these insights allowed us to quantify the binding probability of the whole di- and tripeptide space of proteinogenic amino acids and pin-point the key properties of a strong binder: high hydrophobicity and moderate size of the side-chain in the first position of the peptide.

A common assumption in biochemical studies of transporter proteins is that strong binders are also well transported ^72^. For instance, it was recently demonstrated for three transporters from the MFS, amino acid-polyamine-organocation (APC) and mitochondrial carrier (MCS) superfamily, that stabilizing compounds identified by thermal unfolding assays are also well transported in follow-up radioactive measurements ^72^. In case of DtpB, however, the trend is different, and only mid-affinity peptides (K_D_ between 100 μM and 2.5 mM) are well transported in a reconstituted system. We speculate that this could be a general feature of promiscuous transporters as opposed to highly specific transporters. We also note that transport assays are technically more demanding, so development of high-throughput and robust approaches would help advance our understanding of the mechanisms of transport and their relationship with binding. On the other hand, a combination of *in silico* predictions followed up by selection of potential interactors using a high-throughput assay can effectively narrow down the number of candidates to be tested in a low-throughput transport assay. In case of DtpB, applying such a funnel reduces the list of all possible di- and tripeptides (n = 8400) to a few hundreds of candidates that can be tested for binding by thermal unfolding assays or other techniques. Next, selected peptides in the medium affinity range can be tested for transport using liposome-based assays (pyranine assay or radioactive uptake measurement) ultimately identifying new peptide substrates for DtpB and potentially other promiscuous peptide transporters, including the clinically-relevant human PepT1 and PepT2 transporters. A critical step in this application would be to explore the moiety of the putative ligand that can mimic the N-terminus of a di- or tripeptide and be used for the initial placement of the ligand in the binding pocket. In this work we used a peptide-specific docking protocol, however, a scoring function for small molecules is available in Rosetta ^73^ thus significantly extending the scope of molecules that can be characterized *in silico*.

Our findings also indicate that high affinity peptides believed to be taken up by PepT1 in the human gut, could in fact act as inhibitors, justifying their use as attractive therapies in inflammatory bowel disease (IBD) and colonic cancer ^74–79^. Overall, this work provides a solid molecular and biochemical basis for understanding how structural plasticity of POT’s binding site allows for uptake of a large diversity of ligands. This constitutes a major step forward towards actual structure-based drug design approaches aiming at inhibiting these transport shuttle systems, or at hijacking them to increase drug absorption.

## Acknowledgements

We thank the Sample Preparation and Characterization facility of EMBL Hamburg for support in this project and the beamlines P13 and P14 at EMBL Hamburg for regular access. We acknowledge Rolf Nielsen for initial NanoDSF characterization of DtpB and Instruct-ERIC and the FWO for their support to the Nb discovery and Nele Buys for the technical assistance during nanobody discovery. Funding for the automated crystallography pipelines at EMBL Grenoble was provided by the grants iNEXT (g.n. 653706) and iNEXT Discovery (g.n. 871037) projects funded by the Horizon 2020 program of the European Commission and through Instruct-ERIC. All past and current members of the Löw group are acknowledged for their input to this manuscript. This research was supported through the Maxwell computational resources operated at Deutsches Elektronen-Synchrotron (DESY), Hamburg, Germany. VK and KJ were supported by a research fellowship from the EMBL Interdisciplinary Postdoc (EIPOD) Programme under Marie Curie Cofund Actions MSCA-COFUND-FP (grant agreement number 847543). The authors thank Ulrike Uhrig (EMBL ChemBio facility) for useful discussions on molecular docking.

## Author contributions

Conceptualization: CL, MK, KJ, VK

Investigation: all authors

Software: VK

Writing – Original Draft: VK, CL, MK, KJ

Writing – Review & Editing: all authors

Visualization: VK, MK, KJ

Supervision: CL, MK, KJ, JAM, JS

Funding acquisition: CL, JAM, JS

## Data availability statement

The structures of the DtpB-peptide complexes were deposited to PDB with the following accession codes: 8B18, 8B19, 8B1A, 8B1B, 8B1C, 8B1D, 8B1E, 8B1F, 8B1G, 8B17, 8B1H, 8B1I, 8B1K, 8B1J. Data and code for the pyranine assay, thermal unfolding assay and docking-based prediction of binder probability were deposited to Zenodo with record ID’s 7612027, 7611944, 7612000, and 7586704.

---

**Movie 1: Structural plasticity of the binding pocket of DtpB.** Structural overlay of the 14 different DtpB-peptide complexes. Coordinating residues of the DtpB binding site are shown in sticks (light- and dark-blue for residues of the N- and C-bundle). Peptides are illustrated in sticks and colored white. Initially, a morph of ten DtpB structures bound to dipeptides is presented, but only the dipeptide backbone (white) and coordinating backbone residues of the binding pocket are shown. This is followed by a morph of the DtpB structures bound to tripeptides. Only minor structural changes on the peptide backbone and coordinating residues of the transporter can be observed. This is followed by a morph of all 14 structures and all residues of the binding pocket involved in peptide coordination (including side-chains) are highlighted in light- and dark-blue sticks. The bound peptides are omitted for clarity and the structural plasticity of the binding pocket is visualized. The binding pocket of DtpB can adapt to the different side chains of the peptide, while the coordination of the peptide backbone remains constant.

