## Supplementary file for "Plasticity of the binding pocket in peptide transporters underpins promiscuous substrate recognition"

### Materials and Methods

#### *Gene construction, protein expression and purification of DtpB*

The full-length *dtpb* (Uniprot ID: P36837) gene was amplified from the *Escherichia coli* genome and cloned into a pNIC-CTHF vector [1]. The construct has a C-terminal $His_6$-tag with a TEV cleavage site. DtpB was expressed and purified as previously described [2,3]. Briefly, DtpB was overexpressed in *E. coli* C41(DE3) cells in TB medium. The cell pellet was resuspended in lysis buffer (20 mM NaP, pH 7.5, 300 mM NaCl, 5 % glycerol, 15 mM imidazole) with 5 units/ml DNase I, 1 tablet of cOmplete™ EDTA-free protease inhibitor (Roche)/100 mL buffer, 1 mg/ml lysozyme and 0.5 mM TCEP followed by cell lysis with an emulsifier (EmulsiFlex-C3, Avestin, three passages). The lysate was centrifuged for 10 min at 10,000 g and the supernatant centrifuged for 50 min at 95,000 g (Optima XE-90, Beckman Coulter). The pellet containing the membrane fraction was solubilized in 1 % DDM (Anatrace, sol-grade) for 1 h. The sample was centrifuged for another 50 min at 70,000 g and the supernatant applied to Ni-IMAC beads (ThermoFisher). After 60 min incubation on a rotating wheel, the suspension was transferred to a gravity column. Following two wash steps with lysis buffer supplied with 0.5 mM TCEP, 0.03 % DDM and 30 mM imidazole, DtpB was eluted with lysis buffer containing 0.5 mM TCEP, 0.03 % DDM and 250 mM imidazole. TEV cleavage was performed overnight during dialysis in gel filtration buffer (20 mM HEPES, 150 mM NaCl, 0.5 mM DTT, 5 % glycerol and 0.03 % DDM), and the protein was further purified by negative IMAC. The flow through was then concentrated and loaded on a HiLoad 16/600 Superdex 200 column (GE Healthcare). All steps following cell lysis were performed at 4 °C. The protein was concentrated to 6-8 mg/mL using a 50 kDa cut-off concentrator (Corning Spin-X UF concentrators), flash-frozen and stored at -80 °C until further use.

##### *Selection, expression and purification of nanobodies against DtpB*

41 nanobodies representing 28 different nanobody families were generated against purified DtpB following established protocols [4]. Nanobody constructs contain a C-terminal $His_6$- and EPEA-tag for affinity purification (pMESy4 vector). In total, 30 nanobodies representing the different nanobody families were expressed in *E. coli* WK6 cells and purified following standard procedures [4,5]. In brief, the cell pellet was resuspended in TES buffer (0.2 M TRIS, pH 8, 0.5 mM EDTA, 0.5 M sucrose) supplemented with one protease inhibitor tablet and 5 units/ml DNase. An osmotic shock was performed by the addition of diluted TES buffer to release the periplasmic proteins. The solution was first centrifuged for 20 min at 10,000 g and additionally for 30 min at 40,000 g. The supernatant was applied to CaptureSelect beads (Thermo Fisher Scientific), which were equilibrated with wash buffer (20 mM NaP, pH 7.5, 20 mM NaCl). After ten column volumes of washing, the nanobody was eluted with 20 mM HEPES, pH 7.5, 2 M $MgCl_2$. The nanobodies were further purified on a HiLoad 16/600 Superdex 75 column in 20 mM HEPES, pH 7.5, 150 mM NaCl, 5 % glycerol, concentrated with a 5 kDa cut-off concentrator to 3-10 mg/ml, flash-frozen and stored at -80 °C until further use.

##### *Biolayer interferometry to characterize nanobody binding to DtpB*

The binding of selected nanobodies to DtpB was measured by biolayer interferometry using the Octet RED96 system (Pall ForteBio). The anti-penta-His sensors were incubated in the assay buffer (20 mM HEPES, pH 7.5, 100 mM NaCl, 15 mM imidazole, 5% glycerol, 0.03% DDM) for 10 minutes. Nanobodies were diluted to 5 ug/ml in the assay buffer, purified DtpB was diluted to 200 nM. All experiments were carried out at 22 °C using the following assay parameters: baseline measurement - 180 s, nanobody loading - 300 s, second baseline measurement - 60 s, DtpB association - 300 s, DtpB dissociation - 900 s. Measurements were reference-subtracted and aligned with each other in the Octet Data Analysis HT software v10.0.3.7 (Pall ForteBio), and dissociation constants were determined using a 1:1 binding model.

##### *Crystallization and structure determination of DtpB bound to 14 peptides*

DtpB and Nb132 were mixed in a 1:1.1 molar ratio 1 h prior to crystallization and incubated at 4 °C. Crystallization plates were prepared with an automated liquid handler (Mosquito, TTP Labtech) using the sitting drop vapor diffusion technique with a final drop volume of 300 nL (at 1:1, 1:2 and 2:1 (v/v) ratios of protein to precipitant) and as previously described [6-8]. A set of crystals were automatically mounted and cryo-cooled through the CrystalDirect Method [6,9]. Peptides were purchased from Sigma-Aldrich, Bachem and GL Biochem (Shanghai) and stocks were prepared by weighing the lyophilized powder using an analytical balance and diluting them in ultrapure water. The initial crystals of the ALA bound structure were obtained from the MemGold2 crystallization screen (Molecular Dimensions) at 19 °C. After several rounds of optimization, several DtpB-Nb132 crystals grown in 100 mM HEPES pH 6.7, 30-40 % PEG 400 100-300 mM NaCl, 100-300 mM $MgCl_2$, 10 mM ALA, using a protein : precipitant volume ratios of 1:1, 1:2, and 2:1, diffracted X-rays to 2.5 Å. The structures bound to AF, AI, AL, AQ, AV, NV, SL were obtained from crystals grown in the same conditions as for ALA (100 mM HEPES pH 6.7, 30-40 % PEG 400 100-300 mM NaCl, 100-300 mM $MgCl_2$, and 10 mM of the respective peptide). Structures bound to KV and MS were obtained from crystals grown in 100 mM MES pH 6.0, 30-40 % PEG 400 100-300 mM NaCl, 200 mM $BaCl_2$, and 20 mM of the respective peptides. Structures bound to AFA, AW, AWA were obtained from crystals grown in 100 mM HEPES pH 7.0, 30-40 % PEG 400 100-300 mM NaCl, 200 mM $Li_2SO_4$ and 10 mM of the respective peptide. The APF structure was obtained from crystals grown in 100 mM HEPES pH 7.0, 30-40 % PEG 400 100-300 mM NaCl, 200 mM $Li_2SO_4$ and 4 mM AFA. Here, once the crystal growth seemed appropriate, the crystals were soaked with 25 mM APF and incubated 1 h at 19 °C before harvesting. For all structures, single-crystal monochromatic diffraction datasets of 3600 or 2400 frames were recorded by the rotation method on EIGER 6M detectors with respective oscillation angles of 0.1 ° and 0.15 ° at the P13 and P14 beamlines operated by EMBL-Hamburg at the PETRA III storage ring (DESY, Hamburg, Germany). Collected reflections were indexed, integrated and scaled using the program XDS [10]. The unit cell dimensions were roughly: a = 54 Å, b = 124 Å, c = 169 Å, $\alpha = 90^o$, $\beta = 90^o$, $\gamma = 90^o$ and the space group was identified as P 2 21 21. Initial phases were obtained by molecular replacement using the atomic model of DtpA [3] and a nanobody structure as search models in Phaser as part of CCP4i2 [11] on the ALA bound dataset. The model was further built manually in Coot [12] and refined in PHENIX [13] and Isolde [14] in iterative cycles. Difference map and OMIT map positive peaks clearly indicated the presence of a ligand within the binding site. Fitting the tripeptide ALA in the density led to better agreement between the experimental data and the atomic model. For the other ligand bound datasets, the latter was repeated with the appropriate peptides and also led to better agreement between the experimental data and the atomic models. Visualisation was performed in ChimeraX [15].

##### *Thermal unfolding assay*

The NanoDSF method [16] was used to follow the thermal unfolding event of DtpB with the Prometheus NT.48 device (NanoTemper Technologies, Munich, Germany). Here, the fluorescence at 330 (F330) and 350 (F350) nm are recorded over a temperature gradient scan. DtpB was diluted to 0.35 mg/ml with assay buffer (100 mM HEPES, pH 7.5, 150 mM NaCl, 0.03 % DDM). All ligands were weighed using a fine scale and dissolved in ultrapure water to a final concentration of 50 mM. 18 uL of DtpB at 0.35 mg/ml was mixed with 2 uL of 50 mM ligand/substrate (5 mM final concentration). Samples were incubated for 10 min at room temperature before loading them with standard capillaries into the Prometheus device. The excitation power was set between 15-25 %, and the tested temperature range was from 20 °C to 85 °C. All measurements were done in triplicates and the results were analysed as described below.

#### *Determination of $K_D$ from thermal unfolding curves*

Thermal unfolding data were exported from the manufacturer's software (PR.ThermControl, v. 2.1.2, NanoTemper GmbH), and the F350/F330 ratio data was analysed in MoltenProt (v. 0.1.1) [17] using santoro1988 model to determine melting temperature $T_m$. Problematic unfolding curves, e.g. curves with spikes, were omitted from analysis.

Thermal shifts were calculated by subtracting the average $T_m$ of the apo state ($T_m^{apo}$) and normalizing by $T_m^{apo}$ to obtain $\Delta T_m/T_m^{apo}$. The following equation was used to describe the dependence of $\Delta T_m/T_m^{apo}$ on the peptide concentration (titration curves, [18]):

$$\frac{\Delta T_m}{T_m^{apo}}(L) = \frac{-RT_{std}}{E_{a1}} \cdot ln\left(\frac{K_D}{K_D+L}\right) \quad \text{eq. 1}$$

where L is the total ligand concentration (assuming that it is close to the free ligand concentration [L]); R is the universal gas constant; $T_{std}$ is the standard temperature (298.15 K); $E_{a1}$ is activation energy of unfolding for apo state. The equation was derived from eq. 16 in [18] assuming that the affinity of extra-site binding ($K_{DX}$) is much higher than L.

$\Delta T_m$ and $T_m^{apo}$ are determined experimentally, and since R and $T_{std}$ are constants, a non-linear curve fitting procedure can be used to estimate $K_D$ and $E_{a1}$. According to Hall's model, $E_{a1}$ is a property of the apo state of the protein, so it can be a shared parameter between titration curves of several ligands. Once $E_{a1}$ is known for the protein, the $K_D$ of any new ligand can be estimated from $\Delta T_m$ measured at a single ligand concentration:

$$K_D = \frac{L}{exp\left(\frac{\Delta T_m}{T_m^{apo}} \cdot \frac{E_{a1}}{RT}\right) - 1}$$

In this equation, only $\Delta T_m/T_m^{apo}$ and $E_{a1}$ contain experimental errors, so the uncertainty of $K_D$ can be formulated as:

$$\sigma(K_D) = \frac{\frac{\Delta T_m}{T_m^{apo}} \cdot L \cdot E_{a1} \cdot exp\left(\frac{\frac{\Delta T_m}{T_m^{apo}} \cdot E_{a1}}{R \cdot T_{std}}\right) \cdot \sqrt{\left(\frac{\sigma\left(\frac{\Delta T_m}{T_m^{apo}}\right)}{\frac{\Delta T_m}{T_m^{apo}}}\right)^2 + \left(\frac{\sigma(E_{a1})}{E_{a1}}\right)^2}}{R \cdot T_{std} \cdot \left(exp\left(\frac{\frac{\Delta T_m}{T_m^{apo}} \cdot E_{a1}}{R \cdot T_{std}}\right) - 1\right)^2}$$

We first used eq. 1 to determine the $E_{a1}$ and $K_D$ for three peptides with diverse affinities: AA, AF, AQ. If $E_{a1}$ was not a shared parameter, its value fell into a broad range depending on the peptide (348-479 kJ/mol). Furthermore, the standard deviation for $E_{a1}$ obtained from the covariance matrix of the fit parameters was as high as 25-40 kJ/mol. If $E_{a1}$ was a shared parameter between all three titration curves, a consistent fit could be still achieved with $E_{a1}$ 365 ± 8.2 kJ/mol (standard deviation from the fitting procedure). $E_{a1}$ can be also estimated from a thermal unfolding curve of the apo state by applying a two-state irreversible model. This model was previously derived for differential scanning calorimetry (DSC) and NanoDSF data [17,19]. We fit several unfolding curves of DtpB in absence of any ligand using the "irrev" model implemented in MoltenProt (v. 0.1.1), and in this case the $E_{a1}$ was 371 ± 5.7 kJ/mol, which is in good agreement with the value obtained from ligand titration curves.
Fitting of titration curves was performed using symfit (v. 0.5.3) [20], where $E_{a1}$ was shared between all titration curves, while $K_D$ was specific to each curve. The initial value for $E_{a1}$ was set to 326 kJ/mol (corresponds to the consensus value reported by Hall [18]), and initial value for $K_D$ was 1 μM. Depending on the number of curves in the input, either Nelder-Mead minimization algorithm [21] or differential evolution [22] followed by Nelder-Mead minimization was used. To obtain an independent estimate of $E_{a1}$, the unfolding curves of the apo state were fit using irreversible unfolding model "irrev" implemented in MoltenProt.
Raw and processed data as well as the results of intermediate analysis steps were deposited to Zenodo (https://dx.doi.org/10.5281/zenodo.7612000). Python program for shared-parameter fitting of titration (kd.py) is available from Zenodo (https://dx.doi.org/10.5281/zenodo.7611944).

##### *Determination of $K_D$ using MST*

Peptides were diluted in ultrapure water at a highest possible concentration (usually 50-100 mM), and then 2-fold serial dilutions in water were made. DtpB was diluted to 500 nM in 200 mM HEPES, pH 7.5, 300 mM NaCl, 1 mM TCEP, 10% glycerol, 0.06% DDM. Equal volumes of DtpB and peptide stock were mixed and incubated at room temperature 15-60 minutes. MST was performed using the Monolith NT.LabelFree (NanoTemper GmbH) instrument at room temperature following manufacturer's instructions. Data analysis was performed with MO.Affinity (v. 2.3, NanoTemper GmbH).

##### *Reconstitution of DtpB into liposomes*

DtpB purified in the detergent DM was used for reconstitution into POPE:POPG liposomes *via* the rapid-dilution method. Liposomes were produced by mixing POPE and POPG (Avanti polar lipids) in a 3:1 (w/w) ratio. Chloroform was added to dissolve the lipids and the solvent was removed under vacuum followed by two pentane washes to remove any residual chloroform. The dried lipids were rehydrated in 50 mM potassium phosphate at pH 7.0 (KPi buffer) to reach a final concentration of 20 mg/ml. The lipid mixture was subjected to two freeze-thaw cycles in liquid nitrogen before being stored at -80°C until used. On the day of the reconstitution of DtpB, the appropriate amount of POPE:POPG liposomes were thawed, diluted in 50 mM KPi buffer to a final concentration of 5 mg/ml. Preformed liposomes were obtained by extrusion through a 0.4 µm filter unit (Avanti). DtpB at a concentration of 0.5 mg/ml was added to the preformed liposomes in a 1:60 (w/w) ratio for pyranine assays or in a 1:10 (w/w) ratio for SURFE$^2$R assays. For empty liposomes, the same volume of SEC buffer was added instead. The mixture was left to incubate at 4°C for 1h before rapid-dilution into 60 ml of cold 50 mM KPi buffer and subsequent ultracentrifugation at 100,000 g for 1 hour at 4°C. The liposomes pellet was resuspended in 50 mM KPi buffer and dialysed against 50 mM KPi buffer for 72 h at 4°C with intermitted buffer changes. Dialysed liposomes were harvested for 1h at 60,000 g and 4°C. The liposome pellet was resuspended in 50 mM KPi buffer to reach a final protein concentration of 0.5 mg/ml. Before long-term storage at -80°C, the proteoliposomes were frozen and thawed three-times in liquid nitrogen. Successful reconstitution of DtpB into the liposomes was determined by running the proteoliposomes together with known protein amounts on an SDS-PAGE, followed by densitometry.

##### *Peptide transport measurement using the pyranine assay*

Proton-coupled uptake of peptides by DtpB, reconstituted 1:60 (w/w) in POPE:POPG liposomes, was indirectly measured using the pH-sensitive dye pyranine (trisodium 8-hydroxypyrene-1,3,6-trisulfonate). On the day of the experiment, proteoliposomes were thawed and the appropriate amount taken for the uptake assay. Here, 5 µg of protein per experiment were used. The proteoliposomes were pelleted by ultracentifugation at 100,000 g for 30 min at 4°C and the pellet was resuspended in internal buffer (5 mM HEPES pH 6.8, 120 mM KCl, 2 mM $MgSO_4$) containing 1 mM pyranine. The mixture was subjected to seven freeze-thaw cycles before being extruded 11-times through at 0.4 µm membrane. After that, the proteoliposomes were centrifuged again at 60,000 g for 25 min at 15°C. The supernatant was discarded and the pellet was resuspended in internal buffer. Residual pyranine was removed by applying the resuspended liposomes onto a pre-equilibrated G-25 spin column (Cytiva). The flow-through was collected and the liposomes pelleted at 60,000 g for 25 min at 15°C. The final proteoliposome pellet was resuspended in internal buffer to reach a final volume of 4 µl per experiment.

All peptide stocks were prepared in distilled water and the pH was adjusted so that the peptide at 2.5 mM final concentration in external buffer (5 mM HEPES pH 6.8, 120 mM NaCl, 2 mM $MgSO_4$) was at 7.0. For the pyranine assay, the prepared proteoliposomes were diluted 1:40 into external buffer in a 96-well black chimney plate (Greiner). Fluorescence of pyranine was measured at excitation wavelengths of 415 nm and 460 nm, and an emission wavelength of 510 nm using a Tecan Spark® 20 M Microplate reader. Peptide was added to a final concentration of 2.5 mM and the transport reaction was started *via* the addition of 1 µM valinomycin. Measurements were performed in a minimum of triplicates with individual measurements performed on separate days.

##### *Quantification of transport curves from the pyranine assay*

Data were exported from the plate reader in XLSX format, and the fluorescence signal was calculated by dividing fluorescence at $\lambda_{ex}$=460 nm/$\lambda_{em}$=510 nm by the fluorescence values at $\lambda_{ex}$=415 nm/ $\lambda_{em}$=510 nm and normalised to 1. Normalized fluorescence curves for empty and DtpB liposomes were first checked to identify the time point when the ligand was added. This was done by finding the largest gaps in the time scale. Empty liposome curve was used to determine the slope of the signal drift by linear fit. The drift was subtracted from the DtpB liposome signal to obtain normalized corrected curves ($RFU_{corr}$), which were then fit either with a straight line or with the exponential equation:

$$RFU_{corr}(t) = RFU_{corr}(t_0) - a \cdot \left(1 - e^{-k_t \cdot (t - t_0)}\right)$$

where $RFU_{corr}(t_0)$ is the signal at the starting time point (fixed value), a is the amplitude of the exponent (parameter bounds from 0 to $RFU_{corr}(t_0)$), $\tau$ is the time constant of the observed transport reaction, i.e. time required for the signal to decrease by 1/e of its initial value (parameter bounds from 50 s to infinity), $t_0$ is the starting time of the measurement (parameter bounds from starting time value to ending time value).

Next, the most appropriate model (linear or exponential) for each curve was selected. The residuals of both fits were calculated and assessed for normality using D'Agostino's K-squared test with statistical significance 0.05. Next, the F-statistic was computed for the squared summed residuals (RSS) of both fits:

$$F = \frac{\frac{(RSS_{linear} - RSS_{exp})}{(p_{exp} - p_{linear})}}{\frac{RSS_{exp}}{(n - p_{exp})}}$$

where $RSS_{linear}$, $RSS_{exp}$ – residuals squared and summed for linear or exponential fit, $p_{linear}$, $p_{exp}$ – number of fit parameters (2 parameters in linear fit, 3 parameters in the exponential fit), n – number of data points in a curve. The critical value $F_{crit}$ was computed from F-distribution with 0.01 statistical significance and degrees of freedom ($p_{exp}$ - $p_{linear}$) and (n - $p_{exp}$).

If at least one of the fits had residuals that were not normally distributed, the F-test could not be applied to identify the best model, and the exponential model was selected. If the F-statistic was negative, then the linear model was selected, because the exponent could not fit the curve sufficiently well. If residuals were normally distributed and the F-statistic was above $F_{crit}$, then the exponential model was selected, otherwise the linear model was selected.

For comparison of transport curves from different ligands the instant slope at the starting time point of the measurement slope($t_0$) was computed. In case of exponential fits, this value corresponds to a/τ, while in case of linear fits, slope($t_0$) equals the slope of the line.

All measurements contained a positive transport control using 2.5 mM AA. If the AA curve did not exhibit τ below 150 s and an amplitude above 0.1, the respective batch of liposomes was considered of poor quality and discarded from subsequent data analysis. Finally, the data were normalized to range 0-1 using slope($t_0$) of buffer-only and AA controls according to the formula:

$$slope(t_0)_{rel} = \frac{slope(t_0)^{peptide} - slope(t_0)^{buffer}}{slope(t_0)^{AA} - slope(t_0)^{buffer}}$$

The slope($t_0$)$_{rel}$ from independent experiments (n=2 or 3) was aggregated using robust equivalents of mean and average (median and median absolute deviation (MAD). This step was necessary to avoid effects of occasional outliers. Curve fitting, data manipulation and visualization were performed in Python 3 with the following modules: numpy [23], scipy [24], pandas [25], matplotlib [26]. Data and code were deposited to Zenodo (https://dx.doi.org/10.5281/zenodo.7612027).

To estimate apparent $K_M$ values, the relative transport rates were loaded into GraphPad Prism (version 9.4.1) and fit to the Michaelis-Menten equation. For $K_i$ measurement for AF, the transport rates at variable AF concentration and fixed AA concentration (2.5 mM) were fit to Hill's equation to determine $IC_{50}$, which was then converted to $K_i$ using the Cheng-Prusoff equation. Curve fitting was performed using scipy function curve_fit.

##### *Peptide transport assay using SURFE$^2$R*

DtpB reconstituted in 1:10 (w/w) into POPE:POPG liposomes was used for uptake assays using an SSM-based monitoring system. Proteoliposomes were diluted to 1 mg/ml final lipid concentration in 50 mM potassium phosphate buffer pH 7.0 buffer and briefly sonicated. Of this solution, 10 ul were then applied to a prepared SF-N1 sensor (Nanion Technologies) according the manufacturer's instructions. Each prepared sensor was subjected to an initial quality control as recommended by the manufacturer. Sensors that did not meet the criteria of a capacitance of 15-30 nF and conductance of more than 5 nS were not used for subsequent measurements. To monitor proton-influx due to peptide transport, the sensor was washed for 2 s at 200 µl/s with resting buffer (20 mM HEPES pH 6.8, 120 mM KCl, 2 mM $MgSO_4$) containing either Alanine or Glycine (inactivating condition), depending on the peptide used in the measurement. Transport is initiated by a wash with resting buffer containing the substrate at varying concentrations (activating condition). To restore the sensor for the next measurement, it was rinsed again for 2s at 200 µl/s under inactivating conditions. Measurements were performed using DtpB containing liposomes and empty liposomes in triplicates on a single sensor, respectively. The peak currents for the empty liposomes in presence of respective peptides were subtracted from peak currents of DtpB-containing liposomes and loaded into GraphPad Prism (version 9.4.1) to determine apparent $K_M$ and $v_{max}$.

##### *Flexible docking of di- and tripeptides into DtpB*

Flexible peptide docking was accomplished using the Rosetta FlexPepDock protocol [27] (Rosetta version 3.12-2020.08+release.cb1caba). Initial models of all possible di- and tripeptides were generated using the Rosetta BuildPeptide application and then aligned to the N-terminus of AF from the experimentally determined DtpB-AF complex (without nanobody) using the Biopython PDB module [28,29] (v. 1.78). These models were minimized (relaxed) using the Rosetta relax application [30] and subjected to the FlexPepDock refinement protocol to obtain 200 docked models. GNU parallel [31] (v. 20180922) in combination with Rosetta's internal parallelization were used to enable efficient processing of the data on a computer cluster.

Peptides with known experimental structures (native models) were used to evaluate the docking performance. First, the residues that build up the native DtpB-peptide interface were identified by performing a contact search within 4 Å distance from all non-hydrogen atoms of the peptide. These residues were used to align the docked models to the native model, and the root mean square deviation (RMSD) between the interface residues was recorded (rmsRC). Next, the RMSD between the backbone of the docked and native peptide was calculated (rmsBB_offset). Then the backbone docked peptide was aligned to the backbone of the native peptide and RMSD between the backbone (rmsBB) and side-chains (rmsSC) was calculated. rmsBB and rmsSC characterize the predicted conformation of the peptide; rmsRC characterizes the conformation of DtpB, and rmsBB_offset characterizes the quality of the overall placement of the peptide in the context of DtpB. All alignments and RMSD measurements were done using gemmi [32] (v. 0.4.3). A cutoff for near-native RMSD was set to 1.5 Å. FlexPepDock protocol uses the "reweighted score" to rank docked models, however, it was observed that in many cases the score does not show a consistent correlation with low RMSD values, in particular, rmsBB_offset. Receiver-operator characteristic (ROC) was used to quantify the performance of the "reweighted score" as a binary classifier between hits (rmsBB_offset below or equal 1.5 Å) and non-hits (rmsBB_offset above 1.5 Å) among all peptides with known native structures. The area under curve (AUC) was 0.68 (AUC for a random classifier is 0.5). Since the FlexPepDock application was designed for peptides that are 5-30 residues long, this result may hint that the default default scoring terms (i.e. weights applied to the components of the Rosetta energy function [33]) are not appropriate for DtpB/peptide interaction. This issue can be tackled by reweighing these terms [34]. To accomplish this in a systematic fashion, Rosetta energy terms for all peptides were extracted using per_residue_energies application and then used to train a regularized logistic regression binary classifier (LR) between hits (rmsBB_offset below or equal 1.5 Å) and non-hits (rmsBB_offset above 1.5 Å) using scikit-learn [35] (v. 0.24.2). The robustness of the classifier was checked using group k-fold cross-validation (k=8), corresponding to 2-3 peptides in the test set and 9-10 peptides in the training set. The AUC in this case was 0.79 ± 0.16 (average ± standard deviation). Next, leave-one-out cross-validation (LOOCV) was used to compare the performance LR and "reweighted score". In case of dipeptides LR outperformed "reweighted score" (AUC 0.89 ± 0.08 vs 0.58 ± 0.14, average ± standard deviation), while for tripeptides "reweighted score" showed better performance (AUC 0.74 ± 0.07 vs 0.61 ± 0.2, average ± standard deviation). Thus, for dipeptides the final hits were selected by taking the top 10 models ranked by the probability to be a hit reported by LR, and for tripeptides the top 10 models by reweighted_score were used. Additional code for data manipulation was written in Python (v. 3.7.10) using pandas [25] (v. 1.2.4), numpy [23] (v. 1.20.2) and scipy [24] (v. 1.6.2). Docking results and intermediate steps as well as the code are available from Zenodo (https://dx.doi.org/10.5281/zenodo.7586704).

##### *In silico prediction of binding probability*

Rosetta energy terms were averaged between the 10 selected hits (see previous section) and used as features for the classifier. The peptides with experimentally determined $K_D$ values were split into two classes: binders ($K_D \leq 1$ mM) and non-binders ($K_D > 1$ mM). These features and class assignments were used to train a regularized LR classifier with hyper-parameter tuning using k-fold CV (k=4) as implemented in scikit-learn [35] (v. 0.24.2). The performance of the classifier was assessed using ROC AUC and k-fold CV (k=5), which was repeated 20 times using shuffled data. The AUC from this CV was 0.67 ± 0.04 (average ± standard deviation), and AUC using the full dataset was 0.88. Model trained on the full dataset was then used to predict binding probability (i.e. probability to have $K_D \leq$ 1 mM) for all possible di- and tripeptides. WebLogo [36] (v. 3.7.8) was used to visualize the consensus sequence of di- and tripeptides that have high binding probability (≥0.8) or low binding probability (≤0.2). Data manipulation was performed using Python (v. 3.7.10) with pandas [25] (v. 1.2.4), numpy [23] (v. 1.20.2) and scipy [24] (v. 1.6.2). Code and data are available from Zenodo.

### Supplementary Tables

*Supplementary Table 1: Selected nanobody sequences*

| Nanobody | **Sequence** |
|---|---|
| Nb119 | QVQLVESGGGLVQAGGSLRLSCAASGRTFSNYRMGWFRQAPGQEREFVASISASGGSTDYVDSAKGRFTISRDNAKNTVYLQMNSLKPEDTAVYVCAATVRFGGTLPGHYNSWGQGTQVTVSSHHHHHHEPEA |
| Nb120 | QVQLVESGGGLVQAGGSLRLSCAASGRTFSNYRMGWFRQAPGQEREFVASISGSGGSTTYADSAKGRFTISRDNAKNTVYLQMNSLKPEDTAVYYCAATVRFGGTLPGHYNSWGQGTQVTVSSHHHHHHEPEA |
| Nb121 | QVQLVESGGGLVQAGGSLRLSCAASGRTFSNYLMGWFRQAPGQEREFVASISGSGGSTRYVDSAKGRFTISRDNAKNTVYLQMNSLKPEDTAIYYCAATVRFGGTLPGHYNYWGQGTQVTVSSHHHHHHEPEA |
| Nb122 | QVQLVESGGGLVQAGGSLRLSCAASGRTFTNYRMGWFRQAPGQEREFVASISGSGGNTVYVDSAKGRFTISRDNPKNTIYLQMNSLKPEDTAVYYCAATVRFGGTLPGHYNYWGQGTQVTVSSHHHHHHEPEA |
| Nb123 | QVQLVESGGGLVQAGGSLRLSCAASRRTFSDYRMGWFRQAPGQEREFVAAISGNGGSTYYVDSAKGRFTISRDNAKNTVYLQMNSLKPEDTAVYYCAATLRFGGTLPGHYNYWGQGTQVTVSSHHHHHHEPEA |
| Nb124 | QVQLVESGGGLVQPGGSLRLSCAASGRTFSNYRMGWFRQAPGQEREFVAASGGGGSTYYVDSAKGRFTISRDNAKNTVYLQMNSLKPEDTAVYFCAATLRFGGTLPAHYNYWGQGTQVTVSSHHHHHHEPEA |
| Nb125 | QVQLVESGGGLVQAGGSLRLSCAMSGRTFSNYRMGWFRQAPGQEREFVAGISGSPSSTSYGDSAKGRFTISRDNAKNTVYLQMNSLKPEDTAVYYCAATLRFGGTLPGHYNYWGQGTQVTVSSHHHHHHEPEA |
| Nb126 | QVQLVESGGGLVQAGGSLRLACAASGRTFSNYRMGWFRQAPGQEREFVAAISETGGTTYYADSAKGRFTISRDNAKNVVYLQMNSLKPEDTAVYYCAATFRFGGTLPAHYNYWGQGTQVTVSSHHHHHHEPEA |
| Nb127 | QVQLVESGGGLVQTGGSLKLACAASGLAFSNYIMGWFRQAPGKEREFVAGINWSGGSTNYADSVKGRFTISRDNAKNTLYLQMNSLKPEDTAVYYCAASTRMLLATYPNLYNYWGRGTQVTVSSHHHHHHEPEA |
| Nb128 | QVQLVESGGGLVQPGGSLRLSCAASGFTFSNYIMGWFRQAPGKEREFVAGINWSGGSTNYADSVKGRFTISRDNAKNTLYLQMNSLKPEDTAVYYCAASTRMLLATYPDLYNYWGRGTQVTVSSHHHHHHEPEA |
| Nb129 | QVQLVESGGGLVQAGGSLRLSCAASGRTFSNYDVGWFRQAPGKEREFVVVIRRSVGATYYGDSVKGRFTISRDNAKNTVDLQMNSLKPEDTAIYYCAAVSRLAILPPTTLKDDYRYWGQGTQVTVSSHHHHHHEPEA |
| Nb130 | QVQLVESGGGLVQAGGSLRLSCAASGPTLSNYAVGWFRQAPGKEREFVAGINWSSGLTYKDAVKGRFTISRDAVKDTVYLQMNSLKPEDTAVYYCAARFGGMLPRQPSGYAYWGQGTQVTVSSHHHHHHEPEA |
| Nb131 | QVQLVESGGGLVQAGGSLRLSCAASGPTLSNYAVGWFRQAPGKEREFVAGINWSSGLTYKDAVKGRFTISRDNVKDTVYLQMNSLKPEDTAVYYCAARFGGMLPRQPSGYNYWGQGTQVTVSSHHHHHHEPEA |
| Nb132 | QVQLVESGGGLVQAGGSLRLSCAASGPTLSNYAVGWFRQAPGKEREFVAGINWSSGLRYKDVVKGRFTVSRDNVKDTVYLQMNSLKPEDTAVYYCAARFGGMLPLQPSGYANWGQGTQVTVSSHHHHHHEPEA |
| Nb133 | QVQLVESGGALVQAGGSLRLSCAASGPTLSNYAVGWFRQAPGKEREFVAGINWSSGLTYKDAVKGRFTVSRDNVKDTVYLQMNSLKPEDTAVYYCAARFGGMLPLQPSGYAYWGQGTQVTVSSHHHHHHEPEA |
| Nb134 | QVQLVESGGGLVQAGGSLRLSCAASGATLSNYAVGWFQQAPGKEREFVAGINWSSGLTYKDAVKGRFTISRDNVKDTVYLQMNSLKPEDTAVYYCAARFGGMLPLQPSAYTYWGQGTQVTVSSHHHHHHEPEA |
| Nb135 | QVQLVESGGGLVQAGGSLRLSCAASGPTLSNYAVGWFRQAPGKEREFVAGINWSSGLTYKDAVKGRFTISRDNVKDTVYLQMNSLKPEDTAVYYCASRFGGMVPLHPSGYTYWGQGTQVTVSSHHHHHHEPEA |
| Nb136 | QVQLVESGGGLVQAGGSLRLSCAASGSIFSINAMGWYRQAPGKQRELVAAINRGGSTNYADSVKGRFTISRDNAKNTVYLQMNSLKPEDTAVYYCNADRTIVPGTGLPFQYDYWGQGTQVTVSSHHHHHHEPEA |
| Nb137 | QVQLVESGGGLVQAGGSLRLSCAASGSIFSFNAMGWYRQAPGKQRELVAAINTGGSTNYADSVKGRFTISRDNAKNTVYLQMNSLKPEDTAVYYCNADTVVATHDARPFTYDYWGQGTQVTVSSHHHHHHEPEA |
| Nb138 | QVQLVESGGGLVQAGGSLRLSCAASGSIFSINAMGWYRQAPGKQRELVAAITSGGGTNYADSVKGRFTISRDNAKNTVYLQMNSLKPEDTAVYYCNADFLDSNLSGRTRLYDYWGQGTQVTVSSHHHHHHEPEA |
| Nb139 | QVQLVESGGGLVQPGGSLRLSCAASGSIFSINAMGWYRQAPGKQRELVAAIASGGSTNYADSVKGRFTISRDNAKNTVYLQMNSLKPEDTAVYYCNADLVDITVVAGPPLLRDYWGQGTQVTVSSHHHHHHEPEA |
| Nb140 | QVQLVESGGGLVQPGGSLRLSCAASVSIFSINAMGWYRQAPGKQRELVAAITNGGSTNYADSVKGRFTISRDNAKNTVYLRMNSLKPEDTAVYYCNADIIDYTAIYPTTYDYWGQGTQVTVSSHHHHHHEPEA |
| Nb141 | QVQLVESGGGLVQPGESLRLSCAASGSIFRLNDMAWYRQAPGKQRELVAAISRGGNTNYADSVKGRFTISRDNAKNTVYLQMNSLKPEDTAVYYCNADQLEGGWGTFTVREDYWGQGTQVTVSSHHHHHHEPEA |
| Nb142 | QVQLVESGGGLVQPGGSLRLSCVVSGSWFSINGMGWYRQAPGKQRELVAAITSGGSTNYADSVKGRFTISRDSAKNTVYLQMNSLKPEDTAVYYCNSLSYGLGPKWDWGQGTQVTVSSHHHHHHEPEA |
| Nb143 | QVQLVESGGGLVQAGGSLRLSCVASGRTFSSYRLGWFRQAPGKERDFVAAISRGGSDTYYADSVKGRFTISRDNTKNTIYLQMNSLKPEDTAEYYCAASSQIGYGTTWTGLREYDYWGPGTQVTVSSHHHHHHEPEA |
| Nb144 | QVQLVESGGGLVQAGGSLRLSCAASDSTLSSYIMAWFRQAPGKEREFVATINWSGSGTSYSDSVKGRFTISRDNAKNTVYLQMNSLKAEDTAIYYCAASSRLQQFTAPRLYAYWGQGTQVTVSSHHHHHHEPEA |
| Nb145 | QVQLVESGGGLVQPGGSLRLTCAASGGTFTSYDMAWFRQAPGKEREFVAAISWFGGVTNYADSMKGRFTISRDNAKNTVYLQMNSLKPEDTAVYYCAARRGLRSMSSNEYDYWGQGTQVTVSSHHHHHHEPEA |
| Nb146 | QVQLVESGGGLVQAGGSLRLSCAVSGRALGSINMGWFRQAPGKDREFVAAIMWTGIRTLYADSVKGRFTISRDDAKNTVYLQMNSLKPEDTGVYYCATETHSIGLIRSASEYDYWGQGTQVTVSSHHHHHHEPEA |
| Nb147 | QVQLVESGGGLVQAGGSLRLSCAASGRALGRSNMGWFRQAPGKDREFVAAIMWTGIRTLYADSVKGRFTISRDGAKNTVYLQMNSLKPEDTGVYYCATDTHSIALIRSASEYDYWGQGTQVTVSSHHHHHHEPEA |
| Nb148 | QVQLVESGGGLVQPGGSLRLSCAVSGGTFSSYAMGWFRQAPGKERELVGSISSGGTTYYADSVKGRFTISRENAKDTTSLQMSSLKPEDTAVYYCAAKRPTWYRTSTFDYWGQGTQVTVSSHHHHHHEPEA |

| | |
|---|---|
| Nb149 | QVQLVESGGGLVQAGGSLSLSCAASGRTLTYYTMAWFRQAPGKEREFVATISWSGGTNYADSVKGRFIIS RDSAKNMVNLQMNSLKPEDTAVYYCAAGRRVVPSAPNGTDYWGKGTQVTVSSHHHHHHEPEA |
| Nb150 | QVQLVESGGGLVQAGASLRLSCAVSGRTLSTAGLGWFRQAPGKEREFVAAISSYGTTTYYADSVKGRFTI SRDNAQSTVYLQMNSLKLEDTALYYCAATATNYRNANSYKYWGQGTQVTVSSHHHHHHEPEA |
| Nb151 | QVQLVESGGGLAQAGGSLRLSCVASAGSFSSYAMGWFRQAPGKEREFVAATSQSGVSTRYAPSVTGRFT VSRDNANNAVYLQMSSLKPEDTAVYYCAARWAIGSLLYQSGYNYWGQGTQVTVSSHHHHHHEPEA |
| Nb152 | QVQLVESGGGLVQAGGSLRLSCVASGGTLNNYAMAWFRQAPGKEREFVAAISWSGSSVRYADSVKGRFT ISRDNAKSTEYLQMSSLKPEDTALYYCAARFGFVITTAPTTYNYWGQGTQVTVSSHHHHHHEPEA |
| Nb153 | QVQLVESGGGSVQAGGSLRLSCVLSGRTFSSYTMAWFRQAPGQEREFVAGITYMGGRRYADSVKGRFTIS RDTAKNTAYLQMNTLKPEDTAVYSCAARSGLVTTDLATYTYWGQGTQVTVSSHHHHHHEPEA |
| Nb154 | QVQLVESGGGLVQAGGSRRLSCVASGSTFRNYVMGWFRQAPGKEREFVARIGSNNAATHYSYSVKDRFT ISRDNAANMVYLQMNSLKPEDTAVYYCAADRWGGGTYPFWGQGTQVTVSSHHHHHHEPEA |
| Nb155 | QVQLVESGGGLVQTGDSLRLSCAASGLVFSNYAMGWFRQAPGKEREFVAAISRSASATQFADSVKGRFTI SRDNNKNTVYLKMSSLEPEDTAVYFCAADHKFSGTALITVPYEYEYWGQGTQVTVSSHHHHHHEPEA |
| Nb156 | QVQLVESGGGLVQAGDSLRLSCAASGRTFGSIGWFRQGPGMEREFVATITWIGGTSYADSVKGRFTISRD DAKNMVYLQMNSLKPEDTAVYFCALGRGSKLSKDYTSWGHGTQVTVSSHHHHHHEPEA |
| Nb157 | QVQLVESGGGLVQAGGSLRLSCAASGLTIGTSGMGWFRQAPGKEREFVAAITWIGGRRYADFVQGRITIS RDDTKDTMYLQMNSLKPEDTAIYYCAKSRSGLTSTSVSGYDYWGQGTQVTVSSHHHHHHEPEA |
| Nb158 | QVQLVESGGGLVQAGASLRLACAPSTPTLSRYTMAWFRQAPGTERQYVARLSWSGMTDYADSVKGRFTI SRDDAEKTVYLQMNSLKPEDTAVYYCAATSKTYVFTTVLRTRDTYEYWGQGTQVTVSSHHHHHHEPEA |
| Nb159 | QVQLVESGGGLVQAGSSVRLSCAVSGFSFRDYAMGWFRQAPGKEREFVGAIYWSRGNTRYADSADSVK GRVTIARDNAKNTVYLQVDTLKPEDTAVYFCAATGYLATYTIPATGTSYRYWGQGTQVTVSSHHHHHHE PEA |

*Supplementary Table 2: Affinities of selected nanobodies to DtpB determined by BLI*

| Nanobody | $k_{on}$ (nM s$^{-1}$) × 10$^{-5}$ | $k_{dis}$ (s$^{-1}$) × 10$^{-5}$ | $K_D$ (nM) |
|---|---|---|---|
| Nb119 | 8.4 ± 0.1 | 1.9 ± 0.01 | 22.6 ± 0.3 |
| Nb120 | 10.6 ± 0.2 | 5.1 ± 0.03 | 48.1 ± 1.0 |
| Nb121 | 11.2 ± 0.2 | 3.1 ± 0.01 | 27.7 ± 0.5 |
| Nb128 | 9.3 ± 0.1 | 3.7 ± 0.02 | 39.8 ± 0.5 |
| Nb129 | 10.0 ± 0.2 | 1.3 ± 0.01 | 13.0 ± 0.3 |
| Nb131 | 11.4 ± 0.2 | 2.1 ± 0.01 | 18.4 ± 0.3 |
| Nb132* | 12.8 ± 0.2 | 1.9 ± 0.01 | 14.8 ± 0.2 |
| Nb133 | 8.1 ± 0.1 | 5.7 ± 0.03 | 70.4 ± 0.9 |
| Nb136 | 3.0 ± 0.1 | 2.0 ± 0.01 | 66.7 ± 0.6 |
| Nb137 | 3.5 ± 0.1 | 4.5 ± 0.02 | 128.6 ± 1.2 |
| Nb138 | 1.7 ± 0.1 | 1.9 ± 0.01 | 111.8 ± 2.1 |
| Nb139 | 1.4 ± 0.1 | 1.5 ± 0.01 | 107.1 ± 1.7 |
| Nb140 | 2.9 ± 0.1 | 2.9 ± 0.01 | 100.0 ± 1.1 |
| Nb141 | 5.0 ± 0.1 | 1.3 ± 0.01 | 26.0 ± 0.3 |
| Nb142 | 4.7 ± 0.1 | 2.6 ± 0.03 | 55.3 ± 1.3 |
| Nb144 | 9.1 ± 0.1 | 1.5 ± 0.01 | 16.5 ± 0.2 |
| Nb145 | 5.8 ± 0.1 | 1.5 ± 0.01 | 25.9 ± 0.2 |
| Nb146 | 6.4 ± 0.1 | 2.6 ± 0.01 | 40.6 ± 0.4 |
| Nb148 | 10.6 ± 0.2 | 6.9 ± 0.05 | 65.1 ± 1.3 |
| Nb149 | 6.6 ± 0.1 | 2.5 ± 0.01 | 37.9 ± 0.4 |
| Nb150 | 9.0 ± 0.1 | 2.7 ± 0.01 | 30.0 ± 0.3 |
| Nb151 | 7.6 ± 0.1 | 1.6 ± 0.01 | 21.1 ± 0.3 |
| Nb152 | 8.4 ± 0.1 | 3.3 ± 0.01 | 39.3 ± 0.5 |
| Nb153** | 12.5 ± 0.3 | 10.3 ± 0.14 | 82.4 ± 2.3 |
| Nb154 | 6.4 ± 0.1 | 1.8 ± 0.01 | 28.1 ± 0.3 |
| Nb155 | 5.6 ± 0.2 | 1.6 ± 0.03 | 28.6 ± 1.2 |
| Nb156 | 12.0 ± 0.2 | 2.9 ± 0.01 | 24.2 ± 0.4 |
| Nb157 | 9.3 ± 0.1 | 1.3 ± 0.01 | 14.1 ± 0.2 |
| Nb158 | 11.6 ± 0.2 | 2.8 ± 0.01 | 24.1 ± 0.4 |
| Nb159 | 2.4 ± 0.1 | 1.4 ± 0.01 | 58.3 ± 0.8 |

* selected for crystallization
** poor signal in the BLI assay

*Supplementary Table 3: Experimental validation of predicted binders*

| Peptide | Binder Probability | experimental $K_D$ | Interpretation |
|---|---|---|---|
| EA | 0 | no $T_M$ shift | True negative |
| GPE | 0 | no $T_M$ shift | True negative |
| PY | 0 | 206.7 ± 126.5 μM | False negative |
| YA | 0.01 | 2.4 ± 0.9 mM | True negative |
| YYR | 0.01 | no $T_M$ shift | True negative |
| HH | 0.02 | 4.3 ± 2.5 mM | True negative |
| SH | 0.13 | 352.0 ± 38.1 μM | False negative |
| RGD | 0.52 | 9.1 ± 6.7 mM | False positive* |
| GD | 0.6 | no $T_M$ shift | False positive* |
| AH | 0.67 | 101.2 ± 35.8 μM | True positive |
| AY | 0.88 | 68.7 ± 17.3 μM | True positive |
| LLA | 1 | 277.9 ± 136.1 μM | True positive |

* ambiguous probability value

### Supplementary Figures

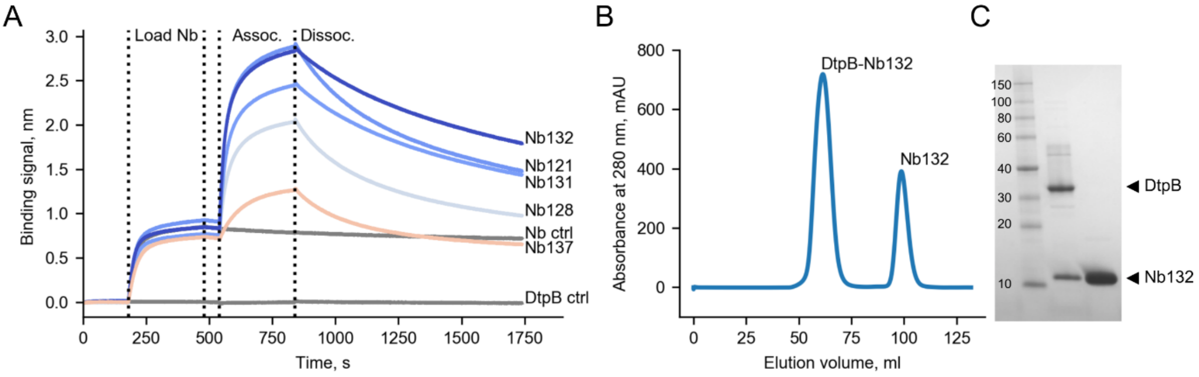


**Supplementary Figure 1: Nanobody Nb132 forms a stable complex with DtpB.** (A) Bilayer interferometry (BLI) was used to assess the binding of several nanobodies raised against DtpB. The two control conditions, where only a nanobody (Nb ctrl), or only DtpB (DtpB ctrl) were loaded on the sensor, are shown in grey. Vertical dashed lines indicate the steps of the assay: (i) loading of the nanobody, (ii) association of the complex, i.e., addition of DtpB, (iii) dissociation of the complex. (B) Representative size exclusion chromatogram (SEC) of the DtpB-Nb132 complex. (C) Representative sodium dodecyl sulfate polyacrylamide gel electrophoresis (SDS-PAGE) of DtpB-Nb132 complex.

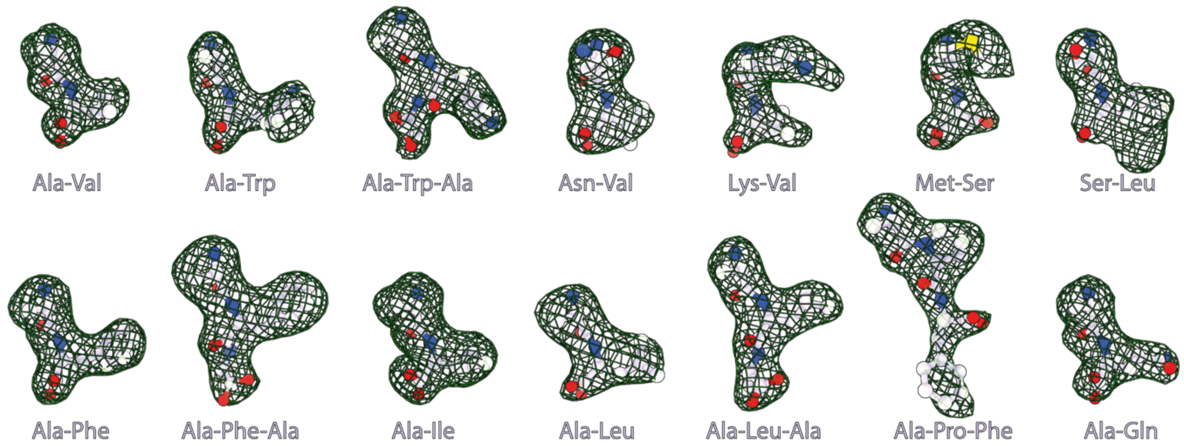


**Supplementary Figure 2: Calculated OMIT maps excluding the 14 modelled peptides**. Polder maps omitting the modelled peptides were calculated for each structure. Positive peaks are displayed as meshed surfaces with a density threshold of +3σ. The omitted peptides are represented as ball and sticks.

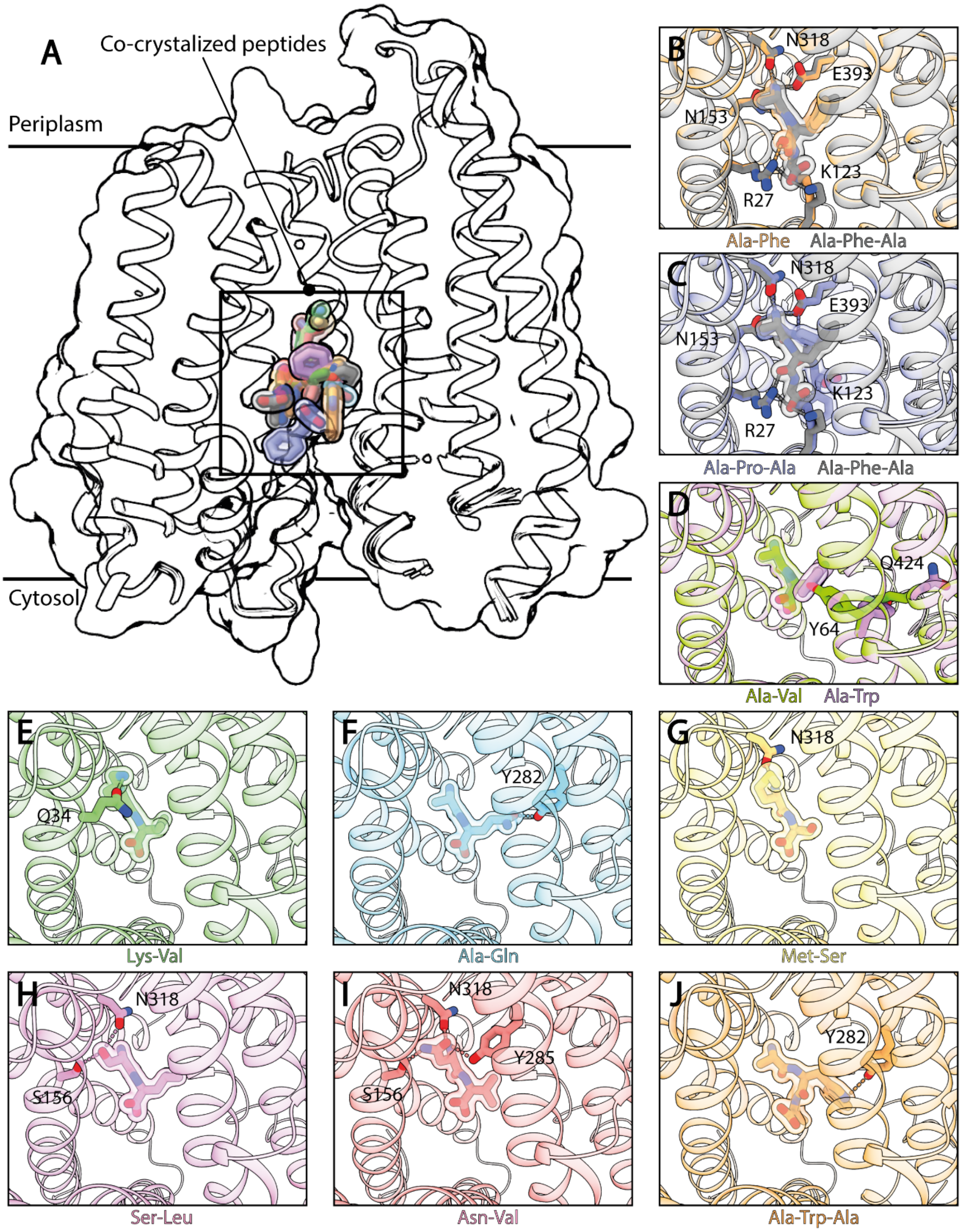


**Supplementary Figure 3: Structural determinants of multi-substrate recognition in DtpB.** (A) The 14 co-crystallized peptides occupy various poses within the binding site. The peptides were coloured differently and the secondary structure elements of the transporter are displayed, as well as the solvent excluded surface. (B, C, D, E) Polar interactions between the side chain of the indicated co-crystallised peptide, and the P1 pocket. (F) Y64 and Q424 rearranges in the presence of peptides with side chains of different bulkiness in the second position (e.g. AV *vs* AW). (G, H) The position of the C-terminus can vary (e.g. in APF *vs* AFA; or AF *vs* AFA), and is accompanied by movements of K123 while the N-terminus remains tightly anchored by the N153, N318, E393 triad. (I, J) Polar interactions between the side chain of the indicated co-crystallised peptide, and the P2 pocket.

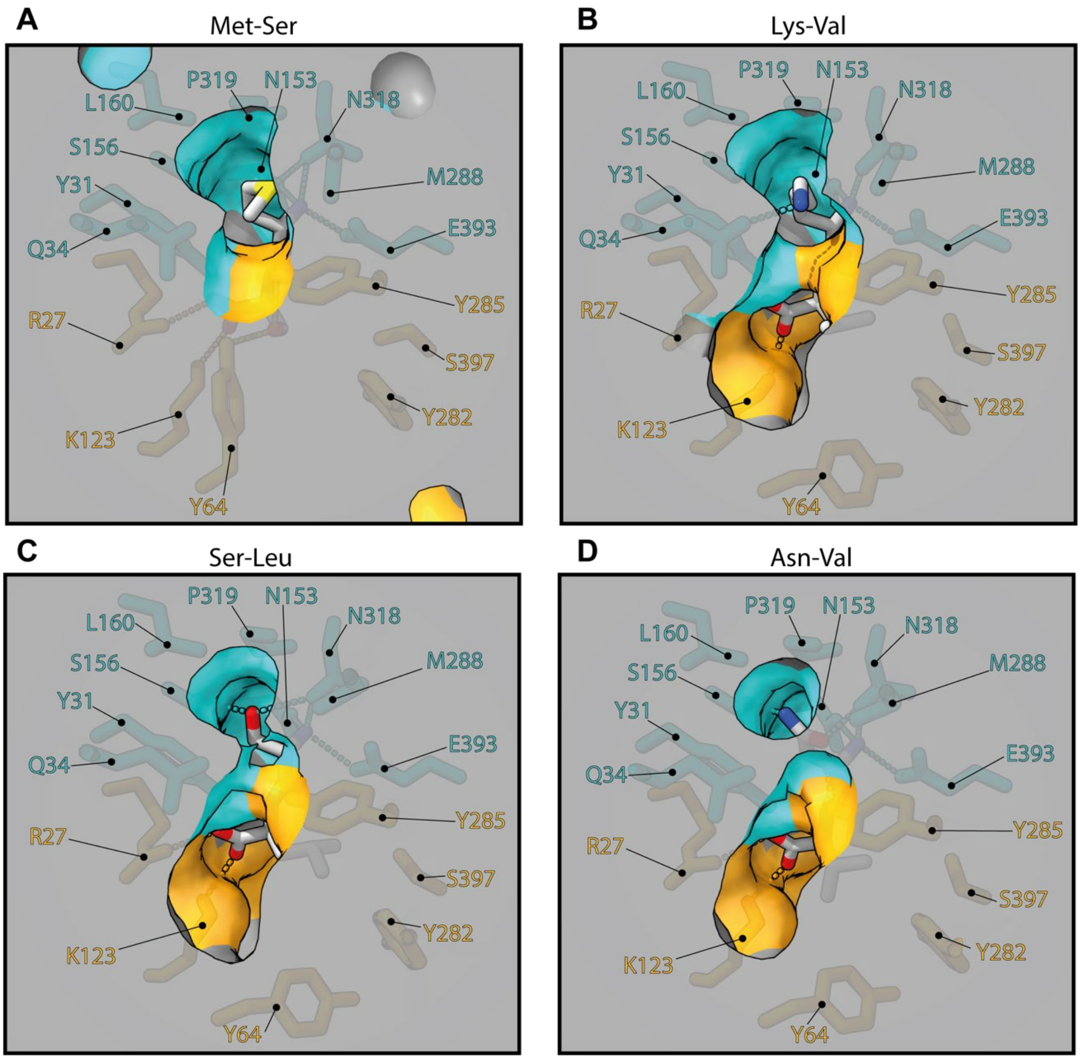


**Supplementary Figure 4: Fitting of side chains in the P1 pocket.** (A) Coordination of the methionine side chain of the MS peptide. (B) Coordination of the lysine side chain of the KV peptide. (C) Coordination of the serine side chain of the SL peptide. (D) Coordination of the asparagine side chain of the NV peptide. Note that P1 remains tight and stable in all these structures.

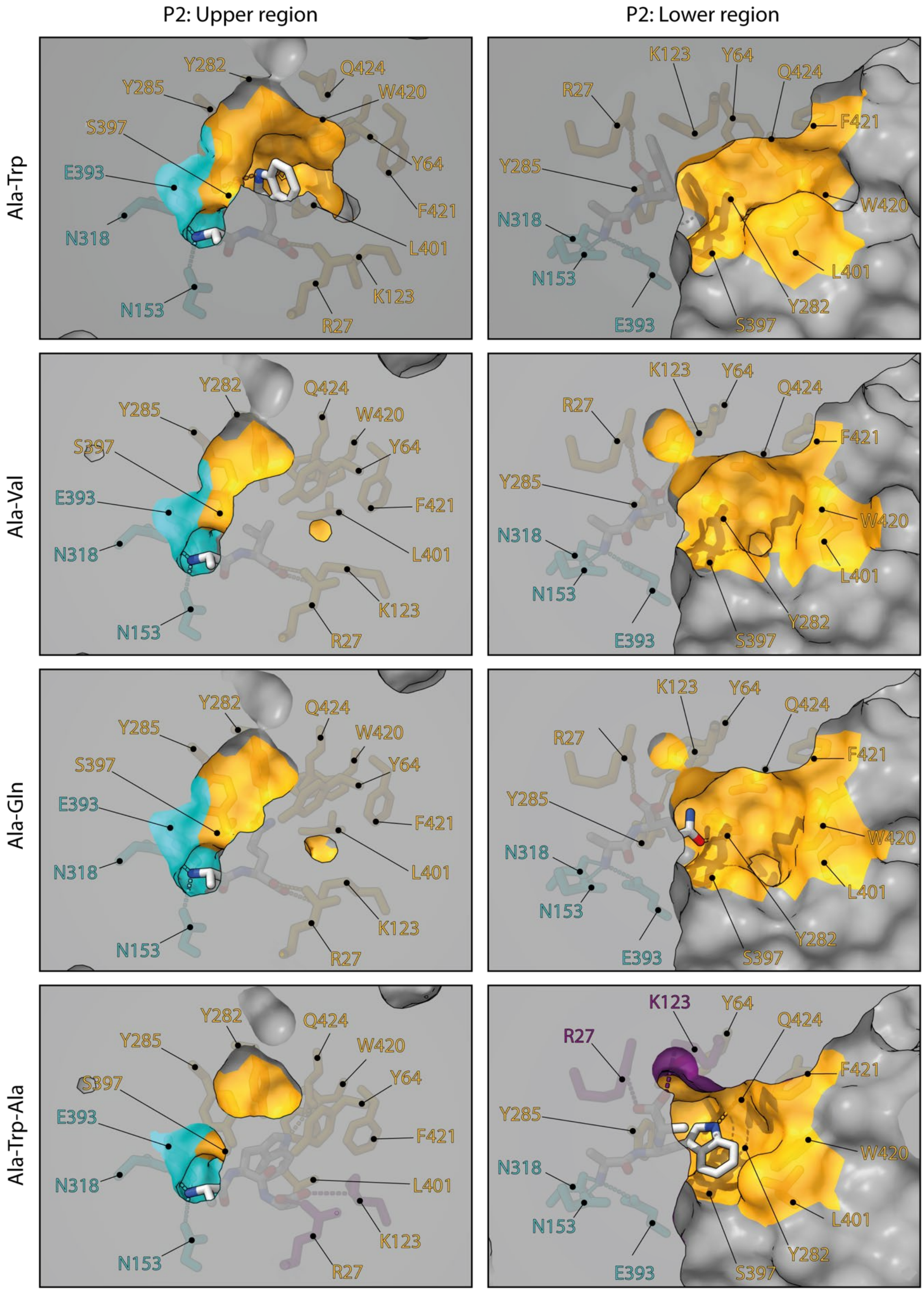


**Supplementary Figure 5: Versatility of the P2 pocket of DtpB.** The upper region of P2 undergoes large rearrangements in the presence of various peptides, by switching the rotamer conformations of Y64 and Q424 (left panels). The indole ring of W2*, in AWA, sits in the lower region of P2, where L401, and W420 can rearrange to cap the cavity.

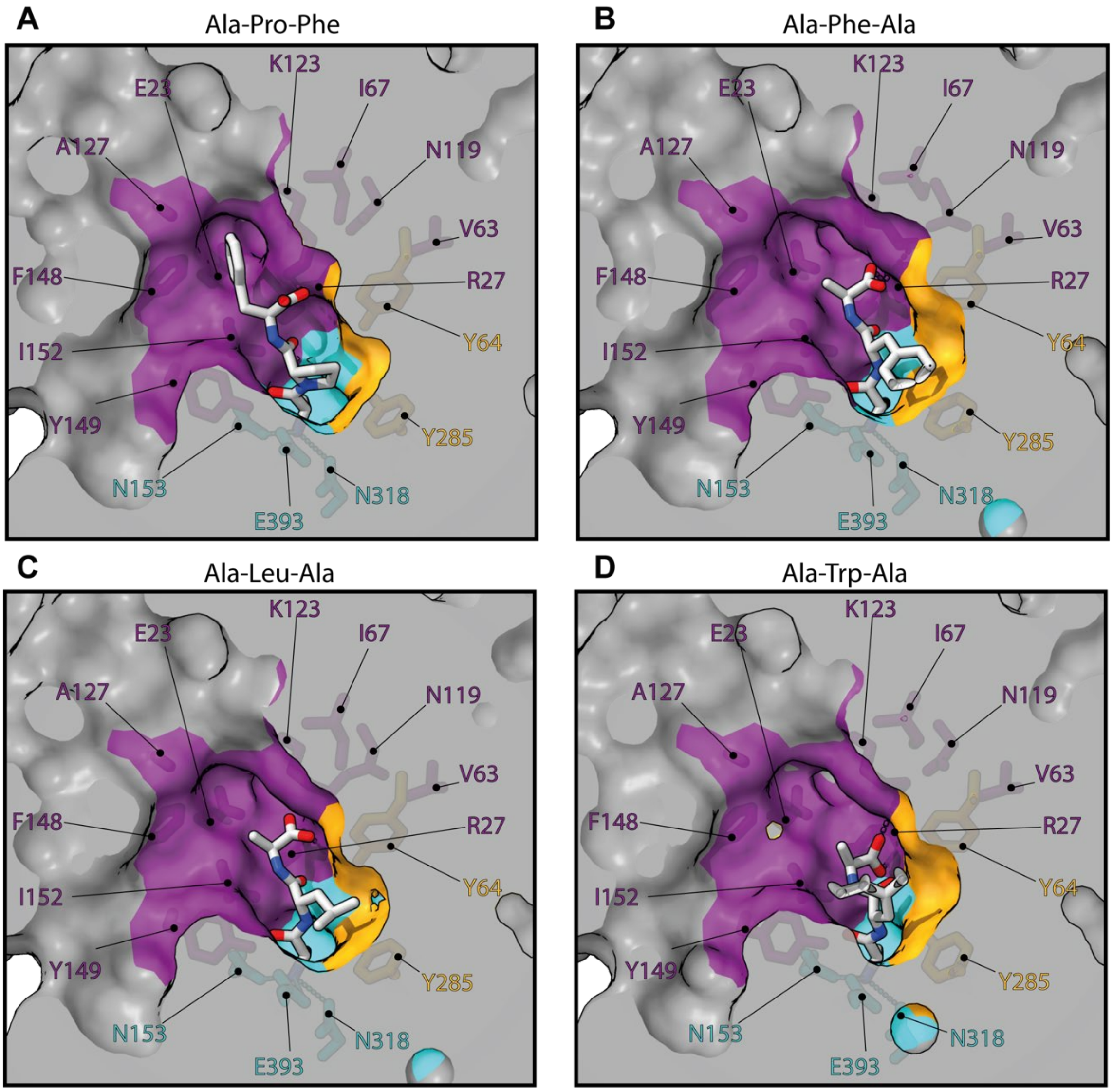


**Supplementary Figure 6: Substrate C-termini and side chains fitting in the P3 pocket.** (A) APF tripeptide, (B) AFA tripeptide, (C) ALA tripeptide, (D) AWA tripeptide. Note that the backbones of the tripeptides can be more extended (A) or kinked (D), and that the C-termini adopt various positions as a result.

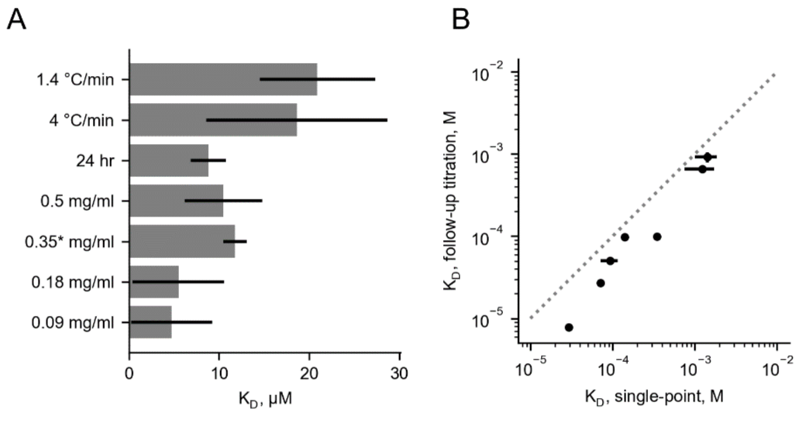


**Supplementary Figure 7: Validation of the approach to determine $K_D$ based on thermal shifts.** (A) Effects of experimental parameters [37] on the estimated $K_D$ of peptide AF: heating rate of 1.4 or 4 °C/min (standard heating rate is 1 °C/min), incubation with ligand for 24 hrs instead of 1 hr, higher or lower concentration of DtpB (the standard value is marked with an asterisk). Error bars denote the standard deviation obtained from the curve fitting procedure. Overall, the calculated $K_D$ for AF is in range 10-20 μM for most conditions. At low protein concentration (0.09-0.18 mg/ml) the signal in the NanoDSF assay is lower, which may result in poor fits and less accurate $T_m$ values. Incubation time does not seem to affect the result, while increased heating rates result in higher $K_D$ estimates with higher uncertainty. Presumably, the increased heating rate decreases the differences in $T_m$, thus affecting the $\Delta T_m$ accuracy, which is important for reliable $K_D$ estimation. (B) Comparison of $K_D$ predicted from single-point $T_m$ measurement at 5 mM peptide concentration (X-axis) with subsequent determination of $K_D$ from a full titration curve (Y-axis). X error bars denote the uncertainty of $K_D$ extrapolation, Y error bars denote the standard deviation obtained from the curve fitting procedure. Dashed line denotes an ideal agreement. We chose peptides (AI, AV, MS, SY, TF, APF, MRF) covering a broad range of $K_D$ (6 – 900 μM), and measured full titration curves with NanoDSF to obtain a more precise estimate of $K_D$. Overall, we observed good agreement, however, the uncertainty increases towards low-affinity ligands. (C) Comparison of $K_D$ determined with NanoDSF and MST. ρ-Spearman's rank order correlation coefficient. Error bars denote the standard deviation obtained from the curve fitting procedure. Dashed line denotes an ideal agreement. $K_D$ estimates from MST have a higher level of uncertainty, and the magnitude of $K_D$ is higher than those predicted from thermal unfolding assays. Importantly, the rank order of the peptide $K_D$ was correct between the two assays (Spearman ρ = 0.76), so the relative affinity is well correlated between MST and thermal unfolding analysis.

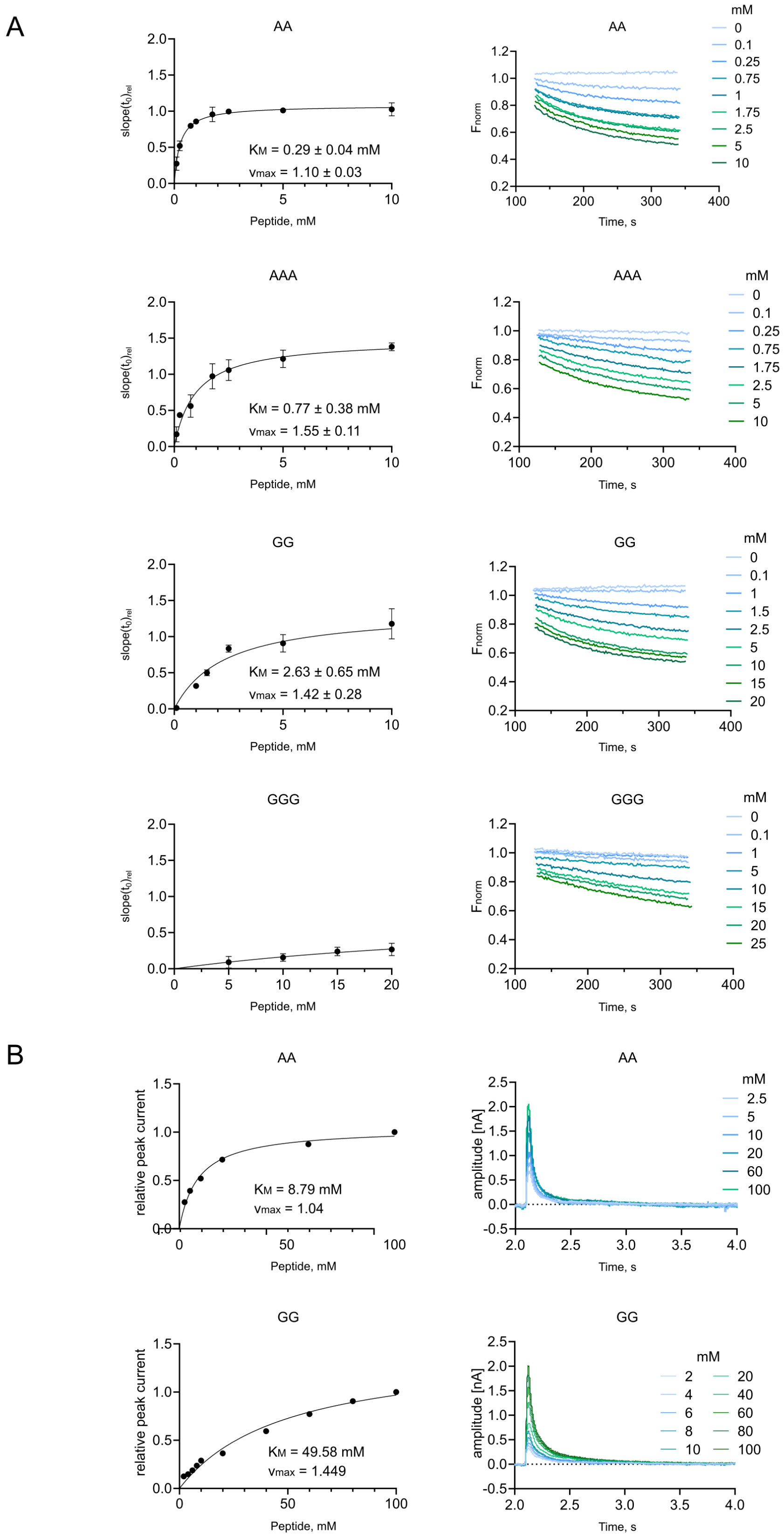


**Supplementary Figure 8: Determination of apparent Michaelis-Menten constant ($K_M$) for model peptides using the pyranine and SURFE²R assay**. (A) Determination of $K_M$ using pyranine assay (left) and exemplary transport curves (right). Error bars denote the standard deviation from n=3

independent measurements. Standard deviation for $K_M$ and $v_{max}$ is obtained from the curve fitting procedure. Insets show raw transport curves. (B) Determination of apparent $K_M$ using SURFE$^2$R for AA and GG (left) and exemplary transport curves (right). In agreement with the $K_M$ estimations from the pyranine assay, the apparent $K_M$ of AA (= 8.79 mM) in SURFE$^2$R is lower than the apparent $K_M$ of GG (= 49.58 mM), i.e. AA is better transported than GG. The differences in the order of magnitude between the derived $K_M$ from pyranine assay (micromolar range) and SURFE$^2$R (millimolar range) are presumably due to the differences in driving forces of the transport. It was previously demonstrated for lactose permease LacY (MFS superfamily) that the apparent $K_M$ decreases several-fold when the membrane potential is applied [38].

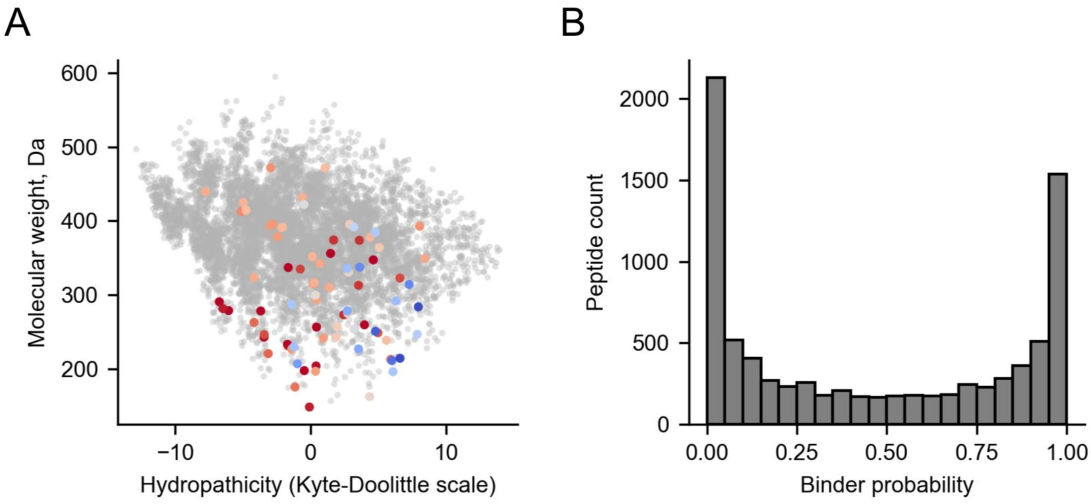


**Supplementary Figure 9: Di- and tripeptide docking into DtpB**. (A) Scatter-plot depicting the space of di- and tripeptides. The X-axis plots hydropathicity of the peptide (Kyte-Doolittle scale) [39], where negative values correspond to polar peptides and positive values indicate hydrophobic peptides. The Y-axis plots the molecular weight of the peptide. Grey points correspond to the peptides that were not studied experimentally in this work. Colored points depict peptides with experimentally determined $K_D$ with high-affinity peptides colored blue and low-affinity peptides colored red. (B) Histogram of the binder probability for all possible di- and tripeptides.

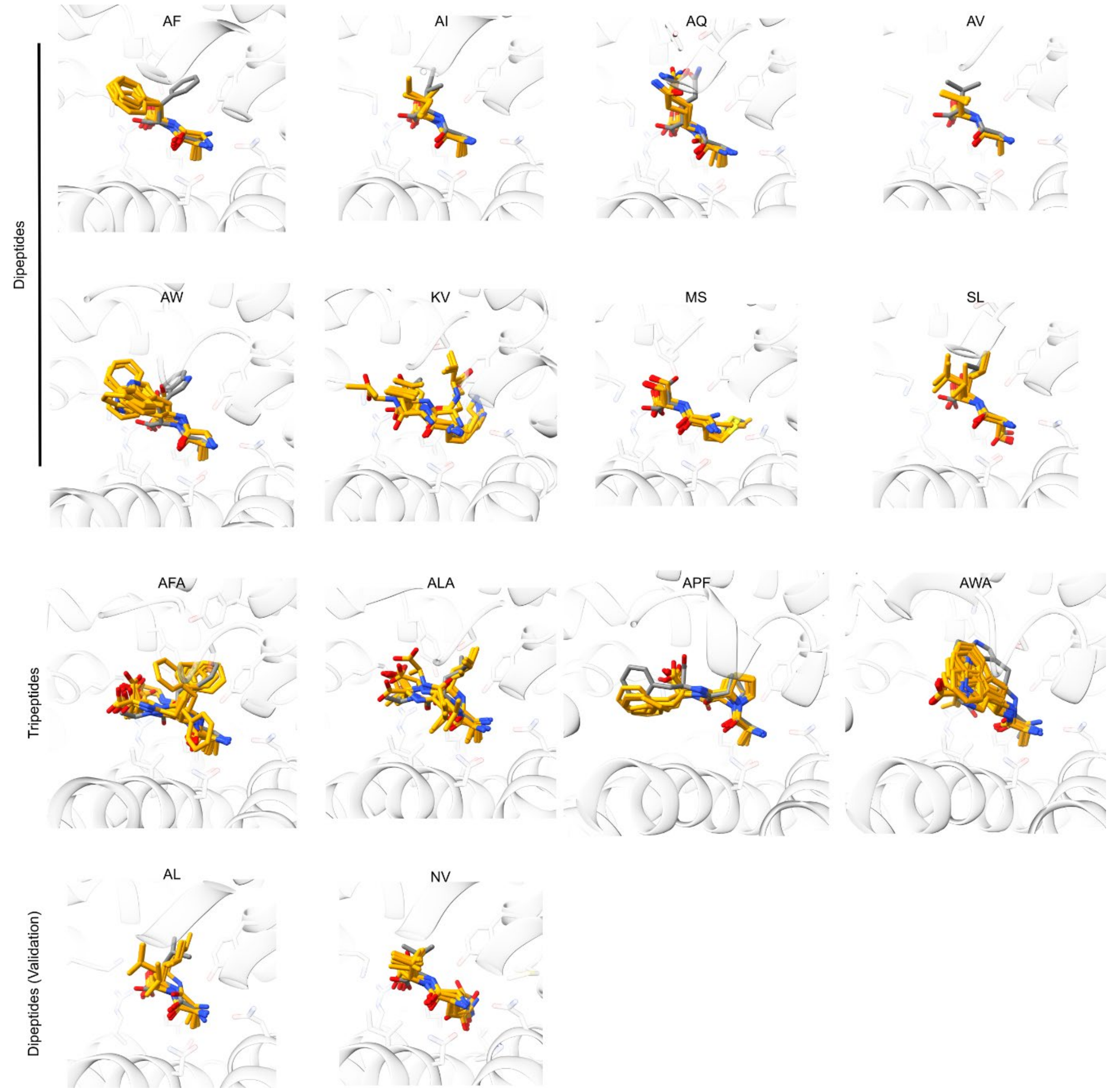


**Supplementary Figure 10: Comparison of experimental and predicted peptide poses.** The binding pocket is viewed from the cytoplasmic side, and only the native structure of DtpB is shown. The native conformation of the peptide is shown in grey, and top 10 docked models are shown in orange.

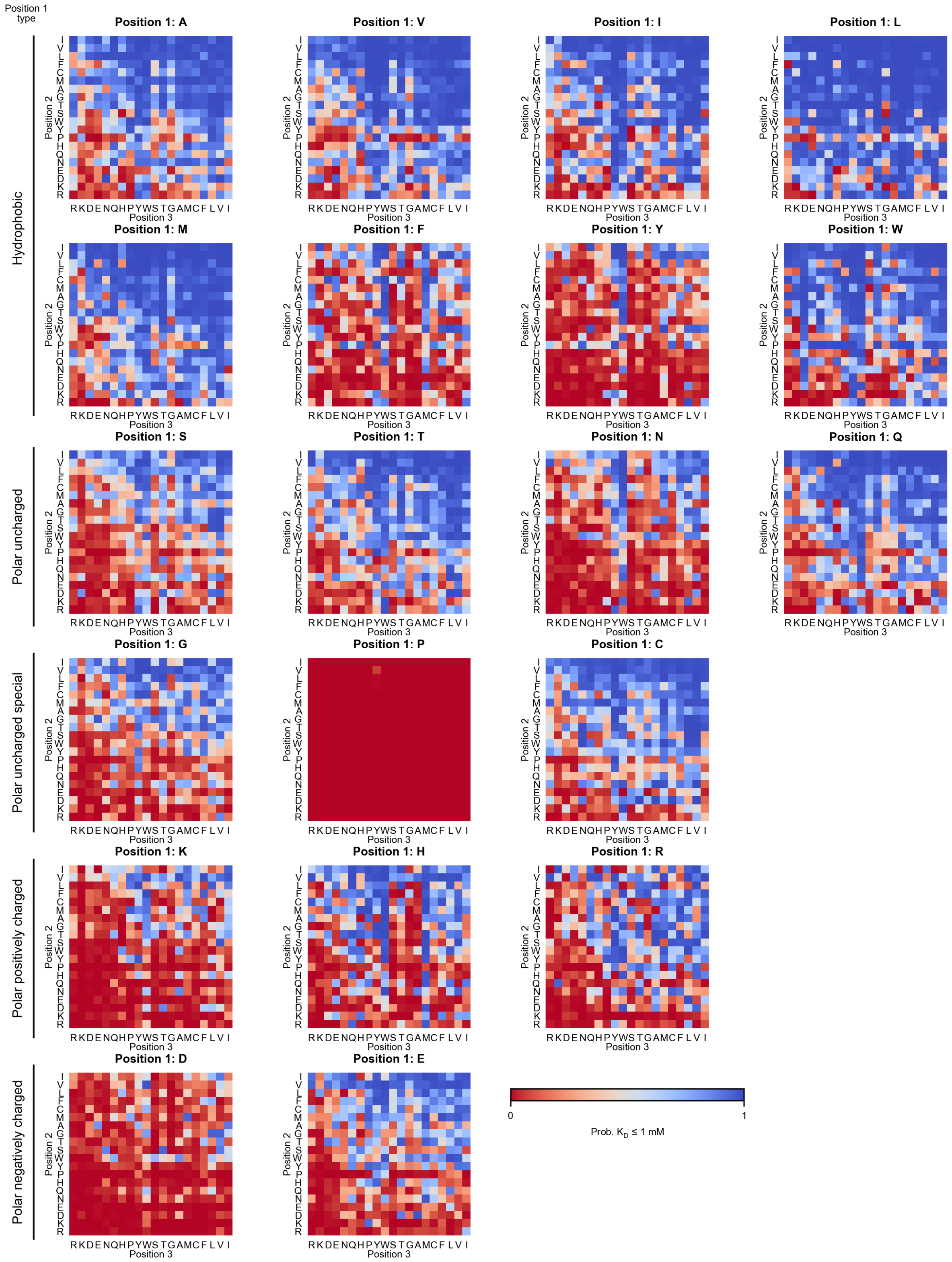


**Supplementary Figure 11: Influence of amino acid identity on the tripeptide binder probability.** Each heat map represents a family of tripeptides with the fixed first position listed in the title of the heat map. Second positions are given in rows and third positions are given in columns. Heat maps are grouped by chemical identity of the first position, and in each group the heat maps are ordered from left to right by increasing molecular weight of the first position. Amino acids in the second and the third position are ordered by Kyte-Doolittle hydrophobicity scale [30] with polar residues on the left/bottom.
